# GTP and lipids control self-assembly and functional promiscuity of Dynamin2 molecular machinery

**DOI:** 10.1101/2021.03.15.435402

**Authors:** Javier Espadas, Rebeca Bocanegra, Juan Manuel Martinez-Galvez, Eneko Largo, Soledad Baños-Mateos, Pedro Arrasate, Julene Ormaetxea Guisasola, Ariana Velasco-Olmo, Javier Vera Lillo, Borja Ibarra, Anna V. Shnyrova, Vadim A. Frolov

## Abstract

Dynamin2 GTPase (Dyn2) is a crucial player in clathrin-mediated endocytosis. Dyn2 is tetrameric in cytoplasm and self-assembles into functional units upon membrane binding. How the curvature activities and functionality of Dyn2 emerge during self-assembly and are regulated by lipids remains unknown. Here we reconstituted the Dyn2 self-assembly process using membrane nanotubes (NT) and vesicles and characterized it using single-molecule fluorescence microscopy, optical tweezers force spectroscopy and cryo-electron microscopy. On NTs, Dyn2 first forms small subhelical oligomers, which are already curvature active and display pronounced curvature sensing properties. Conical lipids and GTP promote their further self-assembly into helical machinery mediating the NT scission. In the presence of large unilamellar vesicles (LUVs), an alternative self-assembly pathway emerges where the subhelical oligomers form membrane tethering complexes mediating LUV-NT binding. Reconstitution of tethering in the LUV system revealed that lipid mixing is controlled by conical lipid species, divalents, GTP, and SH3 binding partners of Dyn2. In membranes with a high content of lipids with negative intrinsic curvature, cryo-EM revealed putative membrane contact sites made by Dyn2 clusters. On such membranes, with GTP lowered to 0.2 mM, both membrane fission and tethering activities become possible, indicating functional promiscuity of Dyn2.

We conclude that GTP and lipids control both extent and topology of Dyn2 functional self-assembly. The function of Dyn2 oligomers evolves from curvature sensing, seen in subhelical Dyn2 oligomers, to curvature creation and fission, seen in Dyn2 helices. Under specific circumstances, such as downregulation of SH3 partners of Dyn2 and GTP depletion, membrane tethering activity can emerge in membrane systems enriched with conical lipids. Hence the Dyn2 functionality is actively adapted to lipidome, explaining its large habitat in the cells and tissues.

## Introduction

Dyn2, the ubiquitously expressed isoform of the "classical" dynamins, is a major player in intracellular membrane trafficking and recycling (Ferguson and De Camilli, 2012; González-Jamett et al., 2013). Recent studies linked Dyn2 to disparate remodeling processes, from mitochondria division to cellular membrane fusion (González-Jamett et al., 2013; Jones et al., 2017; Lee et al., 2016). Yet, the canonical function of Dyn2 is within the clathrin-mediated endocytosis (CME). Analogously to its neuronal homolog Dynamin 1 (Dyn1), Dyn2 is involved in two different CME stages. First, it has a regulatory role during the clathrin-coated pit (CCP) formation and maturation (Bhave et al., 2020). Second, it mediates the scission of the mature CCP from the plasma membrane (Bhave et al., 2020). The scission function of both Dyn1 and Dyn2 is generally ascribed to the protein collar self-assembling on the CCP neck. *In vitro* reconstitution revealed that the collar is a mechano-active helical polymer (Chappie et al., 2011). GTP-driven constriction of the collar produces membrane curvature stress leading to the neck’s fission (Antonny et al., 2016; Schmid and Frolov, 2011). As both classical dynamin isoforms share many similarities, the mechanisms of the helix self-assembly and mechano-action are commonly extrapolated from the Dyn1 helix, which has been more extensively studied. Biochemical and ultra-structural assays revealed that the helical polymerization of Dyn1 is mediated by a set of evolutionarily conserved interfaces residing in the stalk domain of the protein connecting the GTPase (G) head with the membrane-residing pleckstrin homology (PH) domain (Antonny et al., 2016; Schmid and Frolov, 2011). The stalk interactions provide sufficient energy to morph the lipid bilayer into a thin tube fitting into the helical lumen. This way, the Dyn1 helix is believed to emerge on the CCP neck and produce its initial constriction. Significantly, helical self-assembly stimulates the GTPase activity of Dyn1, as the Dyn1 GTPase unit is a G-G dimer (Chappie et al., 2011). The G domains in adjacent helical rungs are set up for cross-dimerization so that the helical self-assembly facilitates the formation of the GTPase units. Cryo-electron microscopy (cryo-EM) showed that the coordinated GTP hydrolysis by multiple G-G dimers in the helix causes robust constriction of the helix and the membrane inside, eventually leading to the scission (Chappie et al., 2011; Kong et al., 2018). It follows that the formation of at least a single helical rung would be required for the G-G dimerization to occur and for the dynamin machinery to function. Indeed, single-molecule fluorescence microscopy analyses of the association stoichiometry of Dyn2-GFP with the CCP revealed that dynamin fission machinery contains 26-30 dynamin molecules, the equivalent of a single helical turn (Cocucci et al., 2014).

The helix to form on a membrane has to overcome the membrane’s elastic resistance to bending. At high concentrations (in the micromolar range), Dyn1 can convert even flat biomimetic lipid membranes into tubes, the process known as membrane tubulation (Liu et al., 2011). However, at lower, more physiologically relevant concentrations, Dyn1 helices only form on highly curved membrane tubes (Roux et al., 2010). Dyn1 binding to these tubes shows steep dependence on the tube’s curvature, with no protein structures formed when the tube radius is greater than 20 nm. Dyn2 relies on the membrane curvature even at high bulk concentrations, only self-assembling into fission-competent structures on curved membrane tubules (Liu et al., 2011). Such membrane curvature "sensing" was suggested to have a distinct meaning in the physiological context. If dynamin fission machinery can only form on sufficiently curved vesicle necks, the neck scission cannot happen prematurely before the CCP is fully mature. However, it is unclear whether extremely short, single-turn Dyn2 helices operating in the cell (Cocucci et al., 2014) have similar curvature creation/sensing capabilities as their microns-long counterparts analyzed *in vitro*. These long helices emerge via a nucleation process from small dynamin oligomers binding to the membrane from the bulk (Roux et al., 2010). The small oligomers could already possess membrane curvature activity and create local curvature comparable with that seen in the long helix. Alternatively, the curvature activity could emerge only upon nucleation of the helical polymerization, when enough oligomers accumulate to bend the membrane into a highly curved tube. A similar dilemma can be envisioned in the cell, where Dyn2 resides in the cytoplasm predominantly as a tetramer (Ross et al., 2011). The minimum number of Dyn2 molecules shown to mediate a vesicle neck’s scission corresponds to that of a single-turn helix (Cocucci et al., 2014), indicating that subhelical Dyn2 oligomers are effective curvature creators. However, their membrane activity has not been directly assessed.

Besides Dyn2 function at the final stage of CME, the curvature activity of subhelical Dyn2 oligomers might underlie its involvement at the earlier stages of the process. Sub- helical oligomers of both, Dyn1 and Dyn2 appear first on low-curved membranes of early CCPs, including the transient non-productive CCPs (Bhave et al., 2020). Recent results show that these small dynamin oligomers actively control CCPs maturation progress, with impairment of their action linked to pathologies (Bhave et al., 2020). Progressive association of sub-helical dynamin oligomers with budding CCPs, seen by fluorescence microscopy (Cocucci et al., 2014; Rappoport et al., 2008; Srinivasan et al., 2018), is in line with their intrinsic membrane curvature sensing capabilities. If they retain the inherent high curvature of the dynamin helix, their local curvature effect should be profound, comparable with that of the clathrin adaptor proteins. As Dyn2 is a hub protein deeply implicated in the CCP proteome, the oligomers’ curvature activity could be an important mechanical factor in the CCP maturation. Furthermore, the oligomers’ curvature-driven sorting towards the emerging vesicle neck might account for the timely self-assembly of the dynamin fission machinery. Finally, the intrinsic curvature activity of small Dyn2 oligomers might also underlie their involvements in the membrane remodeling processes different from endocytosis, such as mitochondria division (Lee et al., 2016) or membrane fusion (González-Jamett et al., 2013). However, *in vitro* analyses of classical dynamins have been mainly focused on complete helical dynamin assemblies implicated in membrane fission. The assessment of smaller oligomers has not been performed, in part, because of the strong dependence of dynamin adsorption on membrane curvature quantified for Dyn1 (Roux et al., 2010). *In vivo* protein partners, such as SNX9 (Srinivasan et al., 2018) and other CCP-associated proteins containing SH3 domain interacting with both Dyn1 and 2, could compensate the low intrinsic affinity of the dynamins for flat and mildly curved membranes (Ferguson and De Camilli, 2012; Yoshida et al., 2004). However, using those auxiliary proteins *in vitro* led to controversial results as their own intrinsic membrane curvature was masking and even competing with that of Dyn2 (Meinecke et al., 2013; Neumann and Schmid, 2013).

Another way to promote dynamin adsorption to slightly curved membranes implicates lipids. Polyunsaturated lipids facilitating the shallow membrane insertion of protein domains (Pinot et al., 2014) also promote the mechano-chemical and membrane remodeling activities of Dyn1 (Pinot et al., 2014). These lipid species are enriched in synaptic membranes, the specific habitat of Dyn1. Dyn2, on the other hand, operates all over the cell and encounters different lipidomes. Yet, there is a unifying lipid element, phosphatidylethanolamine (PE), the conical lipid known to upregulate membrane remodeling. As PE is present in all the membranes known to be remodeled by Dyn2, we reasoned that PE might be the lipid cofactor for Dyn2 activity. That is, PE may control membrane binding of small Dyn2 oligomers and their self-assembly into functional mechano-chemical units.

Following this hypothesis, we reconstituted membrane remodeling activities of small, sub-helical, dynamin oligomers on PE-containing membranes, membrane nanotubes (NT), and vesicles to test this conjecture. Using physiologically relevant amounts of PE, we managed to trap small precursors of Dyn2 helical assembly on highly-curved NTs. These sub-helical Dyn2 oligomers were capable of both sensing and creating local membrane curvature. PE promoted their membrane binding, condensation, and higher-order self-assembly on the NT. On the other hand, GTP drove small precursors’ self-assembly into minimal yet fully functional fission machinery. Surprisingly, we found that these sub-helical Dyn2 units mediated tethering and fusion of membranes containing high PE amounts in the absence of GTP.

Moreover, the same lipid-enhanced tethering activity without GTP was detected for atlastin (Atl), the dynamin family member mediating homotypic membrane fusion (Orso et al., 2009). Thus, while the "traditional" mechano-chemical activity of dynamins is tightly coupled to a well-defined operational specialization (e.g., fusion for Atl and fission for the classical dynamins), the functional promiscuity, seen as lipid-assisted membrane tethering and lipid mixing, emerges with a drop in GTP levels. The observed functional promiscuity at low GTP is further modulated by protein partners’ expression and lipidome alterations, possibly contributing to abnormal membrane morphologies during pathologies related to the dynamin superfamily members (Ali et al., 2019; Zhao et al., 2018).

## Results

### PE regulates functional self-assembly of Dyn2

We begin by testing how PE affects the binding of Dyn2 to LUVs, the template commonly used in functional testing of purified dynamins (Sweitzer and Hinshaw, 1998; Takei et al., 1999). The PH domain of Dyn2 specifically recognizes the highly charged phosphatidylinositol PI(4,5)P_2_ species enriched in the plasma membrane (Klein et al., 1998). Consistently, when LUVs contained 4 mol% of PI(4,5)P_2_, the high amount often used in the *in vitro* assays, Dyn2 binding was virtually unaffected by the PE presence in the membrane (Fig. 1A, S1). However, at lower PI(4,5)P_2_ content (0.5 mol%), the binding showed a steep dependence on PE content, vanishing at 10 mol% of PE (Fig. 1A, S1). We next tested whether the PE presence affects the GTPase activity of Dyn2, known to be stimulated by LUVs (Tuma et al., 1993). Surprisingly, the PE presence was crucial for the stimulation of the GTP hydrolysis even at 4 mol% of PI(4,5)P_2_ (Fig. 1B), indicating that the PE effect goes beyond the stimulation of the protein binding to the lipid bilayer (Pinot et al., 2014). As stimulation of the GTPase activity of dynamins requires helical self-assembly (Sweitzer and Hinshaw, 1998), the much-decreased GTPase activity seen at low PE indicates that PE facilitates LUVs deformations accompanying the formation of Dyn2 helices (Fig. S2). To test this conjecture, we used the Giant Suspended Bilayer (GSB) system recently developed for quantitative analysis of membrane remodeling (Velasco-Olmo et al., 2019). In agreement with earlier observations (Liu et al., 2011), Dyn2 showed limited tubulation activity on GSBs containing a low PE amount even at 4 mol% PI(4,5)P_2_ (Fig. 1C, S3). Yet, the tubulation activity increased gradually with the PE concentration (Fig. S3, 1C), corroborating that PE facilitates helical self-assembly of Dyn2 on low-curved lipid membranes. Hence, PE promotes membrane binding and the helical polymerization of Dyn2, thus controlling the functional self-assembly of the Dyn2 machinery.

**Figure 1.**
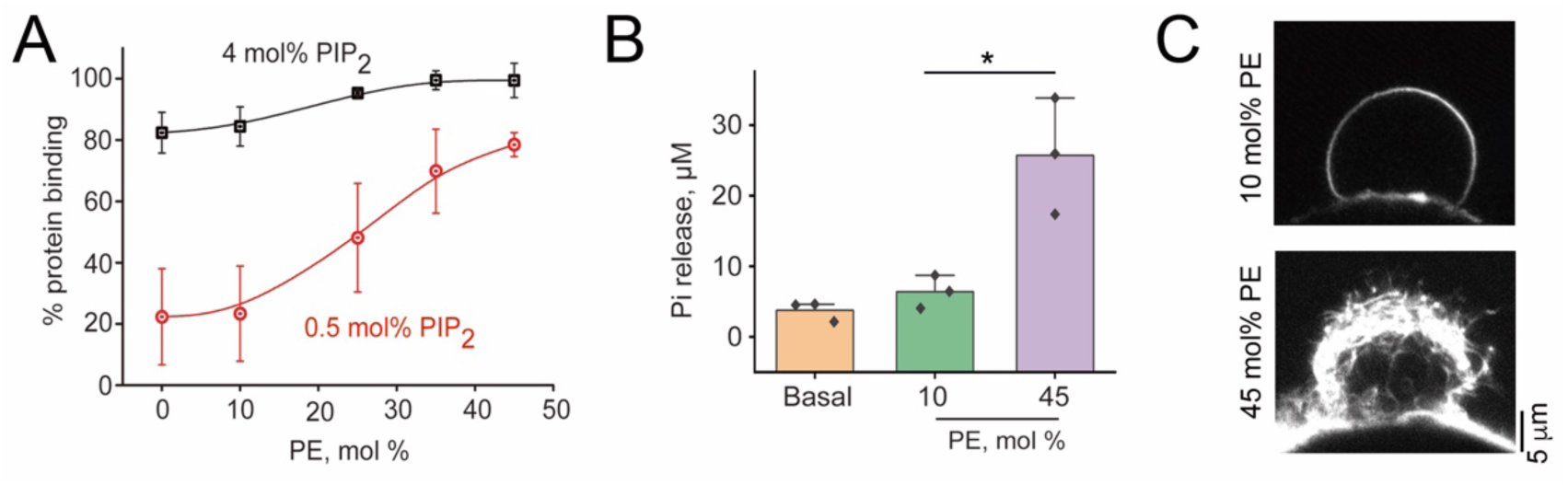
PE enhances membrane binding and helical self-assembly of Dyn2. **A.** Effect of PE on Dyn2 binding to LUVs containing high (4 mol%, black) and low (0.5 mol%, red) amount of PI(4,5)P_2_. See Fig. S1 for the binding quantification details. Error bars are SD. Each data point corresponds to 3 independent experiments. **B.** PE enhances Dyn2 GTPase activity. The graph displays the Pi release measured 30 min after GTP addition in the absence (basal) and presence of 400 nm LUVs containing 4 mol% of PI(4,5)P_2_ and different PE concentrations. Error bars show SD, three independent experiments per condition. Statistical significance: unpaired two-sample *t*-test, \**p* < 0.05. **C.** Representative images of Dyn2-induced tubulation of GSBs containing 4 mol% of PI(4,5)P_2_ in the presence of 10 mol% of PE (left) or 40 mol% of PE (right). See also Fig. S3.

Dyn2 PH domain controls the tubulation activity of Dyn1 and Dyn2 isoforms on low-PE membranes (Liu et al., 2011). Swapping of the PH domains between Dyn1 and 2 was found sufficient to bring the tubulation activity of Dyn2-PHDyn1 chimeric protein to the level characteristic for Dyn1 (Liu et al., 2011). Accordingly, we found that Dyn2- PHDyn1 chimera and Dyn1 effectively tubulate GUVs independently on the PE content in the membrane (Fig. S4). This result suggests that PE affects Dyn2 functionality by modulating the interactions between the PH domain and the lipid bilayer. Specifically, due to its conical molecular shape, PE can facilitate the insertion of the variable loop 1 (VL1) of the PH domain into the membrane core, similarly to polyunsaturated membrane species (Pinot et al., 2014). We note that the augmented membrane insertion has a double effect. Not only it stimulates membrane binding of Dyn2 (Fig. 1A, lower curve), but it also stabilizes the orientation of the PH domain required for the helical self-assembly (Mehrotra et al., 2014) to promote further membrane tubulation (Fig. 1C, lower panel). Conversely, PI(4,5)P_2_ stimulates the binding (Fig. 1A, upper curve), but not the tubulation activity (Fig. 1C, upper panel), so that lipid geometry and charge distinctly regulate the membrane interactions of Dyn2.

The dependence of the tubulation efficiency on the PE concentration in the membrane has a sigmoidal shape characteristic for cooperative processes (Fig. S3). While the tubulation efficiency rises sharply at 25 mol% PE (Fig. S3), the binding efficiency grows gradually (Fig. 1A), suggesting a well-defined nucleation process. We next studied how helical self-assembly follows Dyn2 adsorption to NTs, an established mimic of the CCP neck (Bashkirov et al., 2008; Prévost et al., 2017).

### Dyn2 oligomers sense and create membrane curvature prior to helical self-assembly

We pulled NTs from GSBs, containing 25 mol% PE and doped with a biotinylated lipid species, using a streptavidin-coated polystyrene bead micromanipulated by optical tweezers, as described earlier (Espadas et al., 2019) (see the schematics in Fig. 2A). The GSB membrane reservoir provides moderate tension, setting the NT radius at 32±19 nm (SD, n=85). We further used the optical tweezers to monitor the force acting on the bead along the NT axis in real-time. The axial force measures the tension-driven retraction of the NT to the GSB reservoir. The addition of Dyn2 to an NT, kept at a fixed length, caused the axial force to decrease to close to zero levels (Fig. 2A). The force drop indicated that the Dyn2 helix forming on the NT abolished the NT membrane retraction to the reservoir (Roux et al., 2010). This effect matches the membrane tubulation activity observed for Dyn2 at the same PE concentration (Fig. 1C, S3). In about half of the experiments (29 out of 59 total cases), the force decreased in a bimodal manner, with a clear intermediate stage (Fig. 2A, *F_ads_*), indicating another process that affects the NT mechanics before the helical polymerization happens.

**Figure 2.**
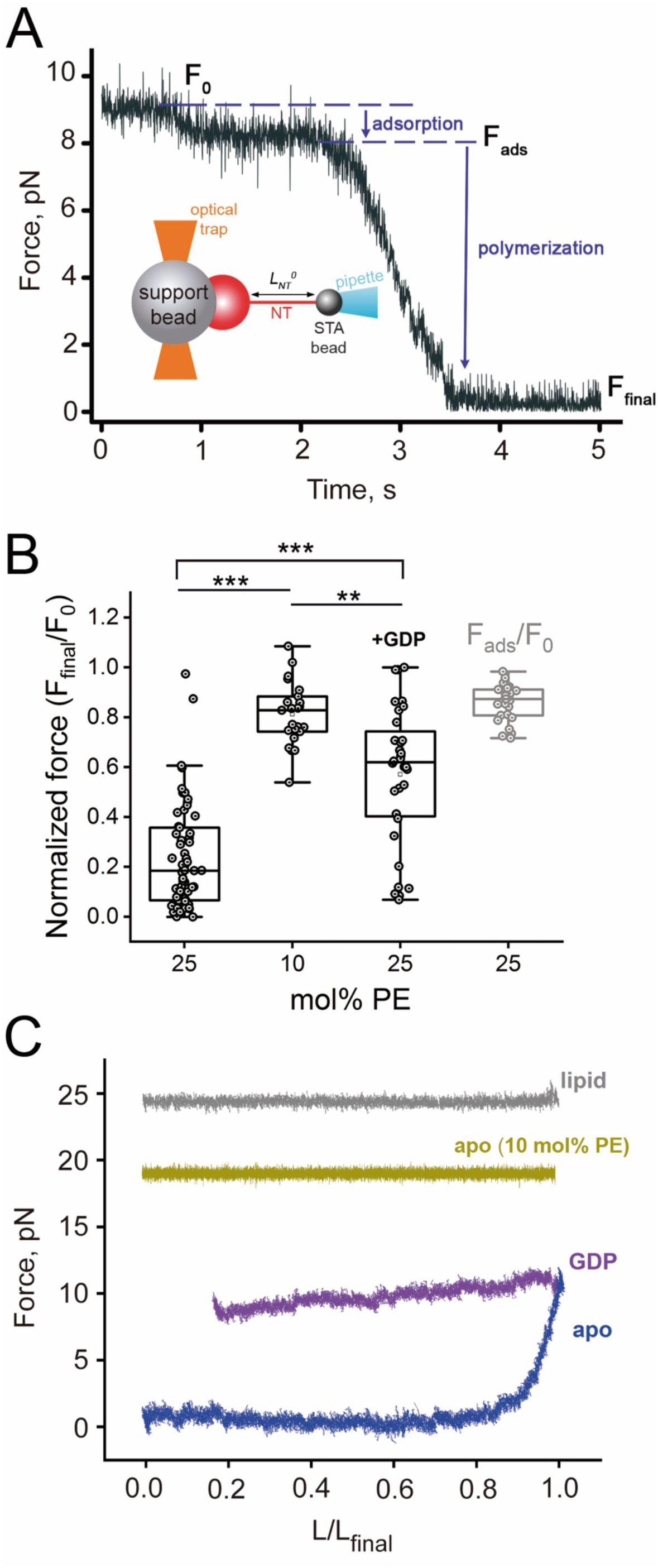
PE affects the mode and the extent of Dyn2-driven stabilization of the lipid NT. **A.** An example of the force measurements experiment. The schematics depict the formation of an NT of length L_NT_^0^ in between a streptavidin (STA) covered bead and the membrane reservoir deposited on the support bead. The graph shows the changes of the force (F_0_) acting on the STA bead along the NT axis (the axial force) upon the addition of 0.5 μM Dyn2 to the bulk. The adsorption of small Dyn2 oligomers on the NT membrane causes the force to decrease to the F_ads_ level. The helical self-assembly followed by the NT coverage by the Dyn2 scaffolds stabilizing the NT lowers the force further to the F_final_ level. The NT membrane contains 2 mol% of PI(4,5)P_2_ and 25 mol% PE. **B.** Boxplot illustrating PE and GDP’s effect on the force decrease measured as in (A). The F_ads_ and F_final_ values normalized to F_0_ are shown. Statistical significance: one-way ANOVA with post hoc Bonferroni test, \*\*\**p* < 0.001, \*\**p* < 0.01. Boxplots show IQR, whiskers indicate the minimum and maximum of the dataset. **C.** The axial force changes upon the NT length decrease with a constant speed *L = L_0_ – (v.t)*, where *t* is the time and *v* = 100 nm/s. Grey trace corresponds to a pure lipid NT, green and blue traces correspond to the NTs decorated by Dyn2 in the absence of nucleotide, and the magenta trace corresponds to the NT retraction in the presence of Dyn2 and 10 mM GDP in solution. The NT membranes contained 2 mol% of PI(4,5)P_2_ and 25 mol% PE, except for the green trace, where the PE content was kept at 10 mol%. Dyn2 concentration was 0.5 μM.

**Figure 3.**
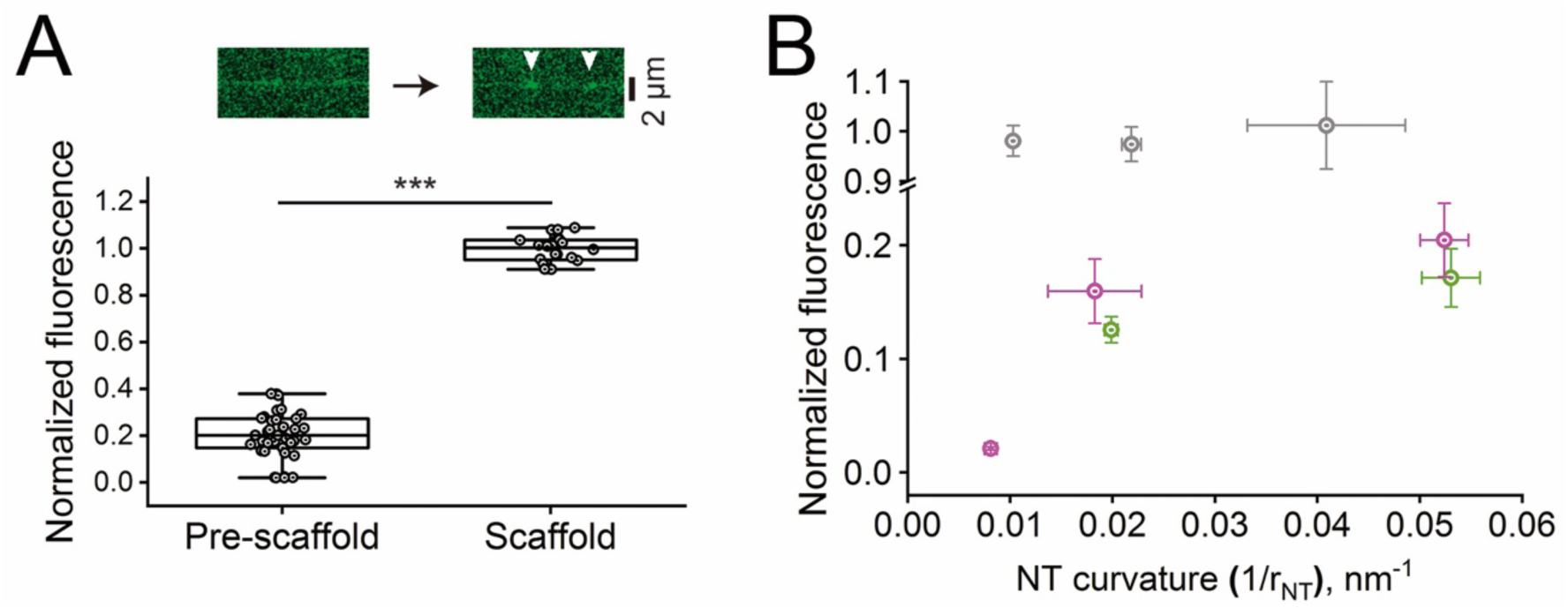
Sub-helical Dyn2 oligomers sense membrane curvature. **A**. The upper panel shows representative snapshots of Dyn2-mEGFP fluorescence distribution over the NT (40 mol% PE) upon the protein addition to the bulk. White arrowheads indicate emerging Dyn2-mEGFP scaffolds. The boxplot shows the fluorescence intensity (per unit NT length, see Fig. S8) of the bound Dyn2-mEGFP in the scaffold regions (Scaffold, n=21) and in between the scaffolds (Pre-scaffold, n=40), normalized to the mean fluorescence measured in the scaffold region. 5 independently obtained NTs were analyzed in each case. Statistical significance: paired-sample *t*-test, \*\*\**p* < 0.001. Boxplots show IQR, whiskers indicate the minimum and maximum of the dataset. **B.** The dependence of the Dyn2-mEGFP surface density in the scaffold regions (grey) and between the scaffolds (magenta) on the NT curvature. The fluorescence density was calculated as the EGFP fluorescence per unit NT length (see panel A and Fig. S8) divided by the NT radius and further normalized to the mean fluorescence density measured in the scaffold regions. The density decreased in the presence of 10 mM GDP, impairing scaffold formation (green). The Dyn2-mEGFP concentration was 0.5 µM in all the cases. Error bars show SD. At least three different NTs were used per each data point.

To gain insight into this process, we used GDP, which is known to interfere with the formation of long-ordered dynamin helices mediating the NT constriction and stabilization against retraction (Bashkirov et al., 2008; Stowell et al., 1999). At 10 mM GDP, the axial force reduction by Dyn2 was diminished almost three times and always happened without a detectable intermediate stage (Fig. 2B, Fig. S5a). Crucially, in 30% of experiments with GDP (n = 27), the force remained constant upon slow reduction of the NT length at a 100 nm/s retraction speed (Fig. 2C, magenta). Such force behavior, similar to the one seen with pure lipid NTs (Fig. 2C, grey), indicates that the NT components, both proteins and lipids, could be returned to the GSB reservoir without creating a barrier for the NT retraction. The partial reduction of the axial force, in this case, could be driven by Dyn2 oligomers too small to constraint the NT retraction and, instead, acting as local curvature creators. Curvature-driven sorting of proteins towards NT is known to produce such an effect (Capraro et al., 2010; Espadas et al., 2019; Sorre et al., 2009). Still, in 70% of experiments with GDP, and in all the cases when the force remained above zero in the absence of nucleotide (n=34), the NT shortening drove the force to the background level (Fig. 2C, blue; S5B, magenta). We associated the force decrease with the condensation of initially dispersed Dyn2 oligomers triggering the formation of a stable protein scaffold.

To further suppress Dyn2 oligomerization, we decreased the PE content in the membrane to 10 mol%. In all trials with such low PE content, we only observed a partial force reduction (n = 24, Fig. 2B, S5C), comparable with that measured at the intermediate stage with 25 mol% of PE (Fig. 2B). Such force reduction indicates concerted action of small Dyn2 oligomers unable to progress to regular helical structures, consistent with the lack of membrane tubulation at low PE (Fig. 1C, S3). Accordingly, the axial force remained constant during the decrease of the NT length in 23 out of 24 experiments with low PE (Fig. 2C, yellow). We conclude that the small Dyn2 oligomers, the precursors of helical self-assembly, act as generic curvature creator units stabilizing the NT against retraction. This activity implies that these oligomers should also be able to sense membrane curvature. To explore their curvature sensing, we employed fluorescence microscopy.

We utilized Dyn2 labeled with monomeric GFP (Dyn2-mEGFP, Methods section), the construct widely used in cellular expression models. We specifically verified that purified Dyn2-mEGFP exhibited GTPase and membrane tubulation activities comparable with wild-type Dyn2 (Fig. S6). As a membrane template for Dyn2-mEGFP, we used multiple parallel NTs produced by the rolling technique from the membrane lamellas utilized in the GSB production (Fig. S7). We used 40% PE to boost helical self-assembly of Dyn2- mEGFP (Fig. S6). The NT radii, 39±15 nm (SD, n=20), were in the range used in the optical tweezers experiments above. Upon addition of Dyn2-mEGFP to the experimental chamber, we first detected the appearance of a weak and uniform mEGFP signal from the NTs, indicating binding of the individual Dyn2-meGFP oligomers (Fig. 3A, upper panel). This initial stage was followed by the fast emergence of brighter mEGFP spots (Fig. 3A, upper panel, arrowheads), indicating the formation of Dyn2 helices (Shnyrova et al., 2013). In between the scaffolds, the mEGFP fluorescence density was on average five times lower (Fig. 3A), in agreement with membrane coverage by dispersed Dyn2-mEGFP oligomers. This two-stage process evokes the dynamics of the axial force decrease (Fig. 2A) though the correspondence is not exact. The helices need to completely cover the NT to prevent the NT retraction to the reservoir and bring the axial force to the background level (Roux et al., 2010) (Fig. 2A, polymerization). The partial force reduction before this moment (Fig. 2A, adsorption) can only be associated with the curvature activity of the non-polymerized fraction of Dyn2 bound to the NT. This fraction is seen in the fluorescence microscopy experiments before the scaffold emergence and then in the regions between the scaffolds (Fig. 3A, S8).

We next assessed whether the Dyn2-mEGFP fluorescence intensity from the scaffolds’ regions shows a systematic dependence on the NT curvature. For that, we normalized the total mEGFP fluorescence to that coming from the lipid fluorescent probe (Rh-DOPE). The mEGFP fluorescence density increased markedly with the NT curvature (Fig. 3B, magenta). In contrast, the Dyn2-mEGFP fluorescence coming from the scaffolds showed no systematic dependence on the NT curvature (Fig. 3B, grey). Hence Dyn2-mEGFP oligomers sense membrane curvature and tend to accumulate in the membrane regions with high positive curvature (Fig. 3B, magenta). This tendency maintains in the presence of GDP (Fig. 3B, green), in agreement with the axial force behavior in the presence of this nucleotide (Fig. 2B, Fig. S5A).

We concluded that the membrane curvature activity of Dyn2 emerges upon its membrane binding before full helical self-assembly and is intrinsic to small protein oligomers. The curvature-driven accumulation of the small oligomers in curved membrane regions, such as NTs or vesicle necks, creates a nucleus for helical self-assembly, thus constituting a plausible mechanism behind the timely emergence of Dyn2 fission machinery in CME. Notably, such functional conversion of small curvature active Dyn2 oligomers to the powerful fission machinery requires moderate PE amounts (e.g., 25 mol%, Fig. 2B). However, higher PE doses further facilitate the curvature of Dyn2 and stimulate other, non-canonical activities of the protein.

### Dyn2 and PE create membrane contact sites mediating membrane tethering and lipid exchange

At 40 mol% of PE, the density of non-helical Dyn2-mEGFP was only five times smaller than that of the helical protein (Fig. 3A). The cryo-EM analysis of Dyn2-driven tubulation of LUVs containing 40 mol% PE revealed that besides well-defined helices (Fig. S2), Dyn2 formed more irregular structures evoking stacking of small protein structures (Fig. 4A, blue arrows (Hinshaw and Schmid, 1995)). The vesicles tubulated by such protein assemblies forming clusters with the vesicles’ lipid bilayers in the cluster often brought into proximity (Fig. 4A, red and white arrows). We relied on fluorescence microscopy to quantify this observed clusterization using the total fluorescence intensity from sub-diffraction spots as a metrics for the cluster size (Jensen et al., 2011). At 0.5 μM Dyn2 the clusterization progressed rapidly and resulted in the formation of massive LUV aggregates (Fig. 4B, C). The clusterization efficiency decreased with the protein concentration, yet scarce clusters could be detected even with a nanomolar concentration of Dyn2 (Fig. S9). The tight contacts between the opposing bilayers, visually associated with tilting of Dyn2 oligomers, indicated bona fide membrane tethering mediated by Dyn2 with a possibility of local merger of the lipid bilayers (Fig. 4A, red arrows). Complete inhibition of the clusterization by the PE removal (Fig. 4B, C) corroborated this hypothesis as PE is known to facilitate the formation of inter-membrane connections and lipid exchange (Chernomordik and Kozlov, 2008; Frolov et al., 2011).

**Figure 4.**
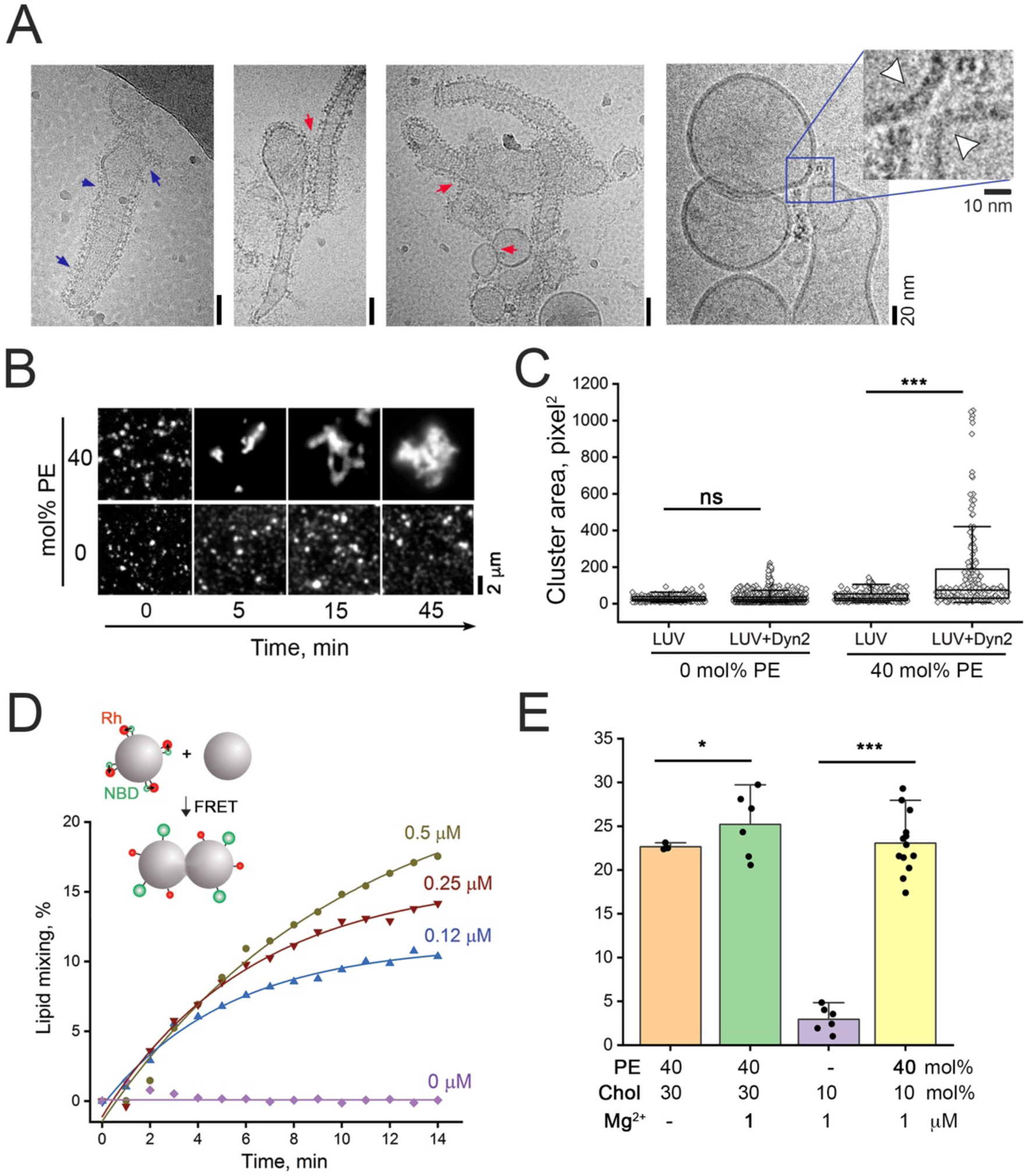
Dyn2-induced tethering and lipid mixing between membranes containing high amounts of PE. **A.** Representative cryo-EM micrographs of Dyn2 assemblies on the membrane containing 40 mol% PE. Irregularities in Dyn2 packing on the membrane (blue arrows) and trans-membrane contacts induced by Dyn2 (red arrows) are seen. The close-up shows the dimple-like structures formed upon Dyn2 tethering of the membranes (white arrowheads). **B.** Time sequence illustrates the clustering effect upon the addition of Dyn2 (0.5 µM, time 0) to LUVs without PE and with 40 mol% PE. Rh-PE fluorescence is shown. **C**. Quantification of the LUV clusterization shown in (**B**). The clusters’ area was measured 10 min after LUVs incubation with Dyn2. The dataset represents at least three independent experiments for each condition. Statistical significance: Wilcoxon signed ranks test, \*\*\**p* < 0.001, *ns* - not significant. Boxplots show IQR, whiskers indicate minimum and maximum of the dataset. **D.** Kinetics of the lipid mixing between LUVs upon addition of different concentrations of Dyn2 as indicated in the graph, measured by the FRET assay (see Methods). The pair Rh-PE and NDB-PE were used as donor-acceptor, respectively (see schematics). NBD-PE fluorescence is shown. **E.** The efficiency of the lipid mixing induced by 0.5 µM Dyn2 at indicated concentrations of conical membrane species (PE and cholesterol) and free Mg^2+^ in solution. The efficiency is measured using the end-point values of the NBD-PE fluorescence increase, as shown in Fig. S13. Statistical significance: paired-sample *t*-test, \*\*\**p* < 0.001, \**p* < 0.05.

To test the conjecture directly, we assessed lipid mixing between the LUVs using Forster Resonance Energy Transfer (FRET) assay (Sekar and Periasamy, 2003). We observed that the Dyn2 addition to LUVs containing 40 mol% PE caused fast lipid mixing (Fig. 4D). The kinetics of the mixing increased with the protein concentration (Fig. 4D), as did the LUV clusterization (Fig. S9). Virtually no lipid mixing was detected without PE (Fig. 4E). These observations confirmed the formation of the inter-membrane bridges mediating lipid exchange in the LUV clusters. As the GTPase activity of Dyn2 relies on Mg^2+^ ion (Vetter and Wittinghofer, 2001), we had the divalent in the protein buffer. We observed that the lipid mixing depended on Mg^2+^ and Ca^2+^ ions but with a more substantial impact of Mg^2+^ (Fig. 4E, S10), suggesting the divalent action on the protein. Accordingly, no lipid mixing was detected at 5 mM of Mg^2+^ without Dyn2 (Fig. S10). Furthermore, the addition of cholesterol, another polyfunctional molecule strongly affecting membrane mechanics, almost completely abolished the requirement for divalent ions (Fig. 4E). Therefore, lipid mixing in this system required the convolution of lipid geometry, electrostatics, and protein-driven tethering, the combo long associated with membrane fusion mediated by specialized protein complexes, such as SNAREs (Jahn et al., 2003; Jensen et al., 2011). Though it was unexpected to see a similar pattern in Dyn2, the protein specialized in membrane fission, membrane tethering activity might not be totally foreign to the protein. Dyn2 belongs to the dynamin superfamily of large GTPases containing both fusion and fission proteins. The functional diversity of dynamins contrasts with substantial structural similarity. However, both fusion and fission activities of dynamins in the cell require GTP hydrolysis.

So far, we considered nucleotide-free Dyn2. We next analyzed how GTP affects the Dyn2 ability to cause membrane tethering and lipid mixing and, in parallel, assessed the effect of PE on atlastin (Alt), the fusogenic dynamin, in the presence and absence of GTP.

### PE-dependent membrane tethering by Dyn2 emerges with GTP depletion, implying functional promiscuity

The addition of GTP (1 mM) promptly inhibited Dyn2-mediated lipid mixing and tethering between LUVs with high PE (40 mol%) (Fig. 5A, B). Notably, the inhibition decreased only gradually with the GTP concentration (Fig. 5B). It follows that at a low physiological concentration of GTP (0.2 mM (Traut, 1994)), both membrane tethering/lipid mixing (Fig. 5B) and membrane fission (Shnyrova et al., 2013) activities of Dyn2 can coexist, implying functional promiscuity of the protein. However, in the cell, Dyn2 operates within the CME proteome and, via its PRD domain, actively interacts with multiple SH3-domain proteins. One of those proteins, amphiphysin, regulates membrane interactions of Dyn2, from helical self-assembly to curvature creation (Ferguson and De Camilli, 2012; Yoshida et al., 2004). We found that this physiological partner of Dyn2 potently inhibits lipid mixing (Fig. 5A). Thus, in the cell, the functional promiscuity of Dyn2 seen *in vitro* is promptly restrained and would be hard to observe under normal circumstances. Yet, in situations associated with depletion of GTP, local or global, Dyn2, in tandem with conical lipids, may become the promoter of membrane tethering and even fusion.

**Figure 5.**
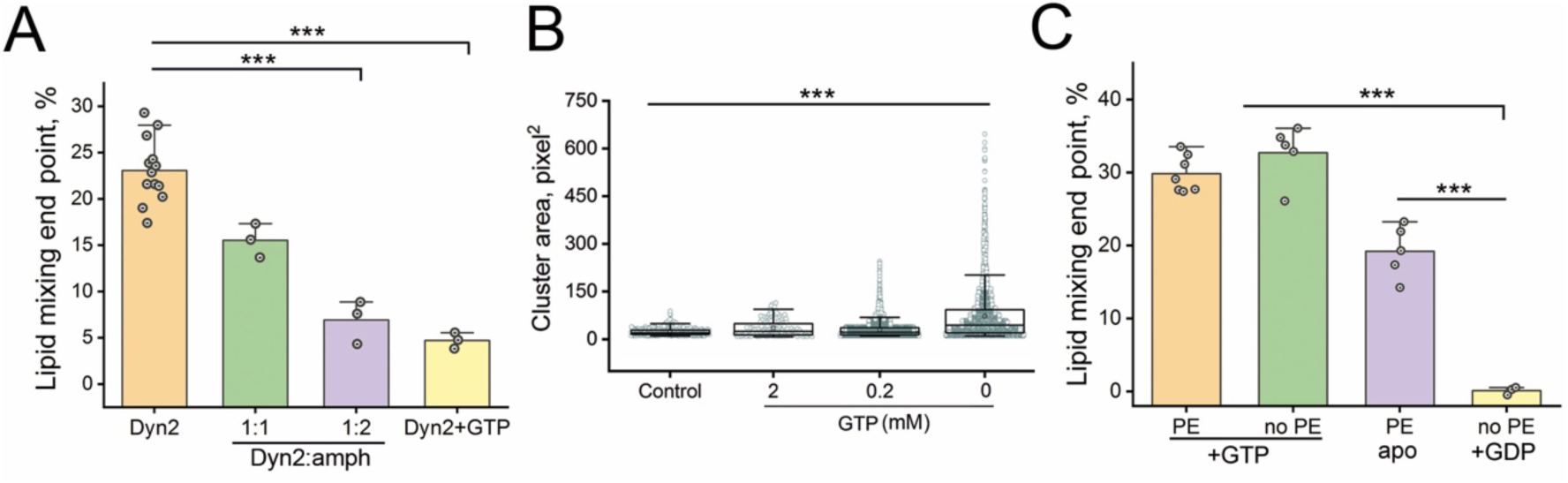
Functional promiscuity of dynamins is controlled by membrane composition, GTP, and protein partners. **A.** The effect of amphiphysin, GTP, and PE on the efficiency of the Dyn2-induced lipid mixing in the LUV system (see Fig. S10); 0.5 µM Dyn2 and 5 mM MgCl_2_ was used in all experiments. Error bars are SD in at least three independent experiments for each condition. Statistical significance: one-way ANOVA with post hoc Bonferroni test, \*\*\**p* < 0.001. **B.** Dependence of LUV clustering (Fig. 4B) on the GTP concentration. Boxplots show IQR, whiskers indicate the minimum and maximum of the dataset. Statistical significance: Kruskal-Wallis ANOVA, \*\*\**p* < 0.001. **C.** Effect of GTP (5 mM) and PE (40 mol%) on the lipid mixing between proteoliposomes containing Atl at 1:400 protein to lipid ratio, in the presence of 5 mM MgCl_2_. Error bars are SD in at least three independent experiments for each condition. Statistical significance: one-way ANOVA with post hoc Bonferroni test, \*\*\**p* < 0.001.

The effect of GTP depletion on Dyn2 interacting with LUVs containing high amounts of PE prompted us to analyze the behavior of Atl in similar circumstances. Atl produces membrane tethering and fusion in the presence of GTP. In the absence of the nucleotide, these activities are known to be abolished (Liu et al., 2015; Pendin et al., 2011; Saini et al., 2014), while maintaining stable tethering does not require GTP (Kim et al., 2017). We reconstituted Atl into LUVs containing 40 mol% PE and no PE and measured lipid mixing in these two proteo-liposome systems. In the presence of GTP, Atl produced effective lipid mixing independently on the PE presence (Fig. 5C), in agreement with the strong fusogenic activity of the protein (Orso et al., 2009). However, as with Dyn2, the presence of PE was critical in the absence of the nucleotide. While no mixing was detected without PE, as expected, Atl showed moderate lipid mixing activity in the presence of PE (Fig. 5C). Hence, both Dyn2 and Atl could promote membrane tethering and lipid exchange between the tethered membrane compartments upon GTP depletion conditions, pointing to dynamins’ ability to form primitive membrane contact sites.

While Atl is designed to mediate local inter-membrane engagement via homo- dimerization (Liu et al., 2015; Pendin et al., 2011; Saini et al., 2014), the membrane tethering activity of Dyn2 at high concentrations was linked to tubulation and protein crowding (Fig. 4A). However, minute LUV clustering detected at much reduced nanomolar concentrations of Dyn2 indicated that the Dyn2 tethering complex might be small. As detection and analysis of scarce tethering events by cryo-EM are hardly feasible, we resorted to single-molecule fluorescence microscopy for stoichiometric analysis of small Dyn2 oligomers forming at extremely low protein concentrations.

### Functional promiscuity of Dyn2 resides in GTP-dependent interaction between small sub-helical Dyn2 oligomers

We used the NT-LUV system to identify the individual tethering events where the LUV attachment to the NT can be easily detected by multicolor fluorescence microscopy. We begin by entrapping individual Dyn2-mEGFP oligomers on the NT containing 40 mol% of PE and doped with 0.1 mol% of Cy5-PE for visibility. We pulse-perfused the NT with a 20 nM solution of the protein. Upon washing off the protein from the bulk, we detected small and mobile Dyn2-mEGFP clusters on the NT (Fig. 6A). Analysis of the fluorescence intensity revealed that the clusters contain around 12 Dyn2-mEGFP molecules (the normalized fluorescence equals 12±6 mEGFP units, Fig. 6B), the amount corresponding to half of the Dyn2 helical turn.

**Figure 6.**
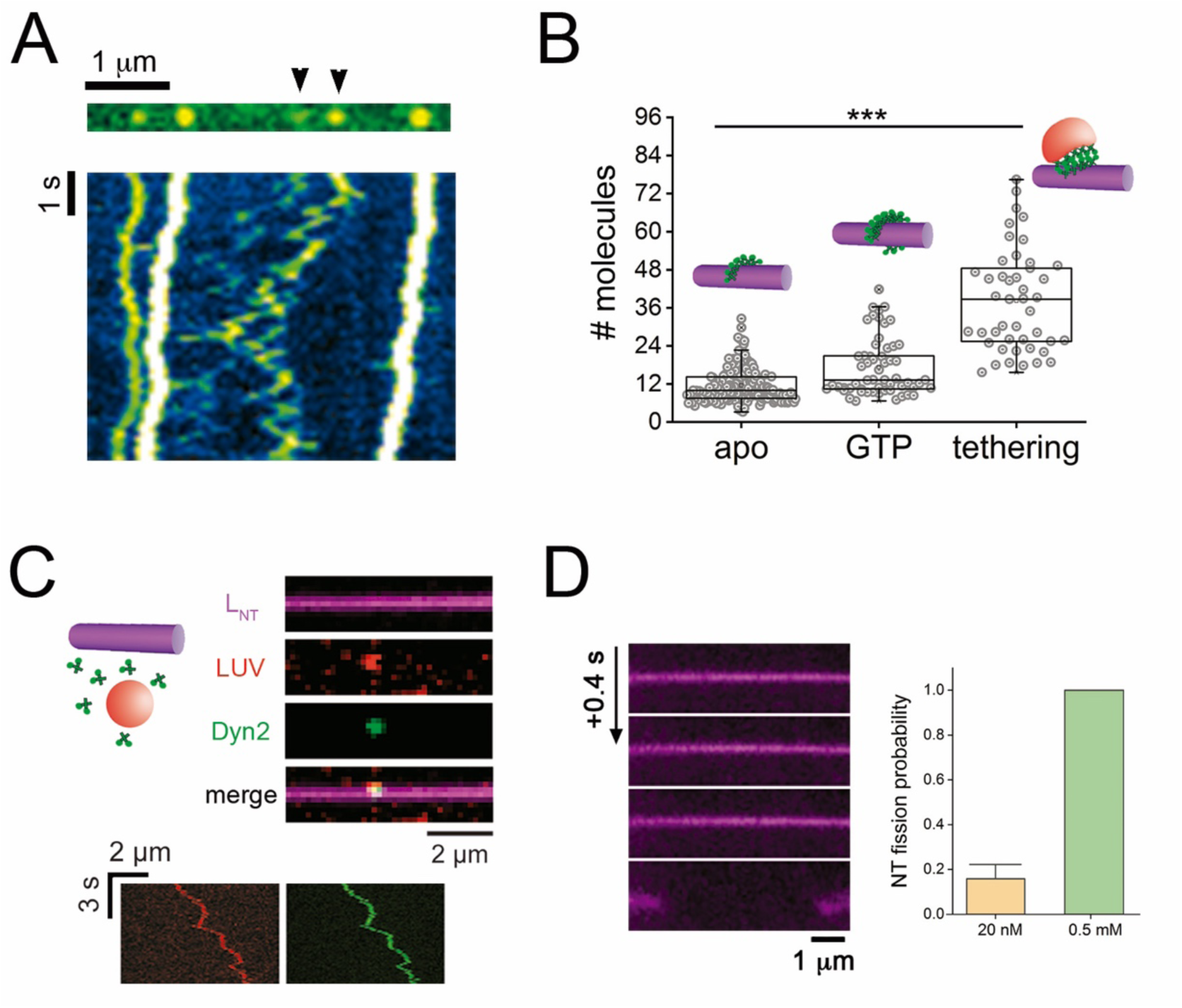
Sub-helical Dyn2 oligomers can form membrane fission and membrane tethering complexes. **A.** The fluorescence micrograph shows Dyn2-mEGFP oligomers bound to an NT. The kymograph illustrates the movement of the oligomers along the NT axis. The pseudo color contrasts the large static oligomers (white) with highly mobile sub-helical oligomers (green, also indicated with black arrowheads in the upper panel). **B.** Number of Dyn2 molecules in the oligomers estimated as total fluorescence intensity divided by the mean intensity of a single mEGFP molecule (see Methods and Fig. S11) in the apo state, upon addition of 2 mM GTP, and upon LUV-NT tethering induced by Dyn2. Boxplots show IQR, whiskers indicate minimum and maximum of the dataset. Statistical significance: Kruskal-Wallis ANOVA, \*\*\**p* < 0.001. **C.** Tethering of 100 nm LUVs (red) to the NT (magenta) by Dyn2 (10-20 nM, green) in the absence of GTP. The kymograph shows how Dyn2 (green) colocalizes with the liposome (red), moving synchronously on the NT axis. **D.** Addition of Dyn2 at low concentration (20 nM) in the presence of 1 mM GTP induces NT fission, albeit with a lower probability than with high protein concentration (0.5 μM). Statistical significance: unpaired two-sample *t*-test, *p* < 0.001.

We next added the protein (20 nM) premixed with LUVs doped with 0.1 mol% of Rh-DOPE fluorophore. Upon washing off unbound material, we detected LUVs sticking to the NTs and colocalizing with Dyn2-mEGFP oligomers, in agreement with the Dyn2 induced tethering between the vesicles and the NT (Fig. 6C). The tethered LUVs moved along the NT axis (Fig. 6C, Movie S1), indicating that the tethering complex is small and not restricting the lateral mobility of the vesicles. Quantification of Dyn2-mEGFP fluorescence colocalized with the moving vesicles showed that the tethering was mediated by Dyn2-mEGFP clusters containing 38±16 molecules (Fig. 6B). The size of the lower group of clusters approached a complete turn of the Dyn2 helix, a structure suitable for stabilizing a lipid neck bridge between the vesicle and the LUV. The structure is about two times bigger than the subhelical Dyn2-mEGFP oligomers forming on the NT without the LUVs (Fig. 6B). This difference in sizes infers that the tethering complex is formed by the interaction between the subhelical oligomer precursors bound to each membrane.

We next explored how these subhelical oligomers react to GTP. With 1mM GTP in the bulk, the size of the Dyn2-mEGFP oligomers bound to the NT increased as compared to the size of the oligomers formed without the nucleotide (Fig. 6B). The larger clusters approached the size previously reported for the Dyn2 fission machinery in vivo (26-30 Dyn2 molecules (Cocucci et al., 2014)). Accordingly, we detected the NT scission events with 20 nM Dyn2, albeit much less frequent than with 0.5 µM of the protein (Fig. 6D, Movie S2).

Together, these experimental observations identify small subhelical Dyn2 oligomers as common precursors of membrane tethering and fission complexes formed by Dyn2. GTP, PE, and protein partners differentially regulate Dyn2 self-assembly along these two pathways, thus controlling the functional promiscuity of Dyn2.

## Discussion

Classical dynamins have the genetically encoded ability to self-assemble into helices in solution (Carr and Hinshaw, 1997) and on curved substrates such as protein filaments (La et al., 2020; Zhang et al., 2020) or membrane necks (Schmid and Frolov, 2011). Helical oligomerization has long been the central subject of Dyn2 research providing the structural and mechanistic foundation for the primary function of Dyn2: the membrane scission. Smaller, subhelical Dyn2 oligomers have escaped scrutiny, as no specific functionality has been ascribed to them, neither in vitro assays nor in the cell. Here we show that subhelical Dyn2 oligomers are curvature-active units that can sense curvature and exercise active, GTP-dependent control on subsequent helical self-assembly. These properties of small Dyn2 oligomers revealed here in the reconstitution assays could provide a new rationale for the appearance and function of small Dyn2 oligomers at the earliest stages of CME. Small Dyn2 units have long been detected in emergent CCPs, as well as caveolae (Bhave et al., 2020; Cocucci et al., 2014)). The protein amounts detected at these early stages of the vesicle budding were generally small, comparable to that constituting the few helical turns of the Dyn2 fission machinery (Bhave et al., 2020; Cocucci et al., 2014). As CCP is also much smaller than the intrinsic curvature of Dyn2 helix, the dynamins oligomers associated with the emergent CCP are most likely small, sub-helical units resembling those detected in the NT in our experiments (see Fig. 6). Curvature-driven sorting of these oligomers towards developing vesicle neck, followed by nucleation of helical self-assembly on the neck, would constitute a plausible mechanism behind the timely appearance of Dyn2 fission machinery at the late stages of the CCP maturation. In turn, the local curvature activity of small Dyn2 oligomers distributed over the CCP could contribute to the mechanical regulation of the CCP budding, thus controlling the maturation process (Bhave et al., 2020). In this context, the small oligomers’ local curvature activity could be amplified by interaction with curvature-generating SH3 partners of Dyn2, the clathrin adaptor proteins (Meinecke et al., 2013).

We further revealed that the size of the small Dyn2 oligomers increases in the presence of GTP (Fig. 6). This observation implies that the oligomers’ curvature activity could be coupled to the GTP hydrolysis cycle, that is, actively controlled. Though we have not directly assessed the GTP-induced oligomerization mechanism, the earlier observations suggest the involvement of nucleotide-driven G-G dimerization. Dynamins have long been known to oligomerize via GTPase domains when arrested in the transition state of the GTPase cycle (Carr and Hinshaw, 1997; Hinshaw and Schmid, 1995). Recent structural analyses revealed that in the presence of GTP, Dyn1 self-assembles into a 2- start helix where the two helical strands interact via the GTPase domains (Kong et al., 2018). This helical arrangement, but no the 1-start helix forming in the absence of GTP, underlies the classical dynamins’ mechano-chemical activity (Kong et al., 2018). We speculate that in the presence of GTP, the sub-helical Dyn2 oligomers can interact laterally, forming a part of a 2-start Dyn2 helix. Whether such mechano-active units could develop on low-curved membranes, such as emergent CCPs, remains to be determined. The curvature and potential mechano-activity of small Dyn2 oligomers could be crucial for the intracellular activities of Dyn2 out of the CME realm. An example of such a process is the control of fusion pore dynamics in various cellular contexts (González-Jamett et al., 2013). It was shown that a Dyn2 octamer controls the expansion of the fusion pore during the HIV entry (Jones et al., 2017). The curved molecules facilitated the expansion, alleviating the membrane bending stress in the pore rim (Chernomordik and Kozlov, 2003), and subhelical Dyn2 arcs would fit for the task. The oligomers’ GTP-driven activation and assembly/disassembly cycles add further versatility to this hypothetic functionality of Dyn2 (Anantharam et al., 2011).

Yet, our experiments show that even in the absence of GTP, the small Dyn2 oligomers could be functionally diverse, causing membrane tethering and even lipid mixing (Figs. 4, 5). Though our observations are limited to in vitro models, we note that the experimental conditions explored here are not unrealistic in the physiological context. Purified Dyn2 oligomers induce membrane tethering already at 0.2 mM of GTP (Fig. 5B), comparable with the intracellular GTP concentration (Traut, 1994). Furthermore, depletion of intracellular GTP and ATP was linked to mutations in phosphoribosylpyrophosphate synthetase 1 (PRS1) associated with Charcot-Marie-Tooth (CMT), the pathology long connected to Dyn2 malfunction (González-Jamett et al., 2014). CMT was also linked with mutations in the Dyn2 pleckstrin-homology (PH) domain (Züchner et al., 2005), which interfere with the GTPase activity of the protein (Muhlberg and Schmid, 2000). Interestingly, such mutations induce vesicular aggregation colocalized with Dyn2 signal in the perinuclear region, indicating that the impairment of the Dyn2 GTPase activity may cause membrane tethering. Yet, under normal circumstances, the tethering activity of Dyn2 would be severely suppressed by its SH3 partners (Fig. 5A (Takeda et al., 2018)), pointing to the critical role of the proline-rich domain (PRD) emergence in classical dynamins evolution.

The functional promiscuity of Dyn2 seen in our reconstitution models points out the importance of the lipidome in regulating both the maintenance and remodeling of intracellular membrane systems. One of the major players there are the cone-shaped lipid species, such as PE or diacylglycerol, imposing negative membrane curvature (Frolov et al., 2011). While unable to make a stable lipid bilayer alone, PE is ubiquitously present in intracellular membrane systems (van Meer, 2005) and specifically enriched in the endoplasmic reticulum and the mitochondrion (van Meer and de Kroon, 2011). As shown here and in multiple earlier studies (Chernomordik and Kozlov, 2003), PE role might converge to making lipid bilayer more susceptible to the action of proteins implicated in membrane remodeling, both membrane fusion (Frolov et al., 2011) and fission (Espadas et al., 2019). The activity of some members of the dynamin superfamily has been linked explicitly to PE. For example, the transmembrane Sey1p, the yeast Atl orthologue, presents a PE-dependent fusogenic activity (Sugiura and Mima, 2016). Though it remains mainly unclear how PE specifically affects dynamins and Dyn2 in particular, our study identifies two possible directions. First, PE might facilitate the membrane insertion of the protein’s PH domain (Ramachandran and Schmid, 2008). The improved anchoring facilitates the protein binding (see Fig. 1A). It also positions the PH domain away from the protein’s stalk domain, thus exposing the interface critical for helical self-assembly and, therefore, the mechano-chemical activity of Dyn2 (Faelber et al., 2011; Ford et al., 2011). Second, PE might facilitate membrane deformations by the Dyn2 via increasing the lipid bilayer compliance to deformation during helical self-assembly (Bashkirov et al., 2020; Shi and Baumgart, 2015).

The cooperation between dynamins and conical lipids creates hotspots for inter- membrane interactions, tethering, and fusion. This synergism and critical involvement of lipid electrostatics (Fig. 4D, S10) are seemingly universal for proteins implicated in intracellular membrane fusion (Jahn et al., 2003; Zimmerberg et al., 2006). But for Dyn2, the dedicated "fission" dynamin, the membrane tethering activity transpires in the absence, or under depletion, of GTP, i.e., under non-physiological or pathological situation(s).

Interestingly, a similar trend is seen in other members of the large dynamin superfamily. In bacteria, soluble protein dynA was shown to mediate membrane tethering and even lipid and content mixing independently on the presence of nucleotides *in vitro* (Bürmann et al., 2011; Guo and Bramkamp, 2019). Other examples are the heterotypic interactions of soluble Cj-DLP1 and Cj-DLP2 from *Campylobacter jejuni* mediating membrane tethering in a GTP-dependent fashion (Liu et al., 2018). In eukaryotes, dynamin-like s- OPA1, the cleavage product of the L-OPA1 proteolysis lacking the transmembrane domain (Ishihara et al., 2006), mediates mitochondrial membrane fission in a GTP-dependent manner (Anand et al., 2014). However, s-OPA also collaborates with L-OPA1 mediating inner mitochondrial membrane fusion independently of its GTPase activity (Ban et al., 2017). While s-OPA1 does not lead to membrane fusion alone, it enhances L- OPA1 fusogenic activity (Ban et al., 2017), likely through membrane tethering. Furthermore, cross-membrane dimerization of L-OPA1 results in membrane tethering in mitochondrial cristae, leading to stabilization of cristae topology. Importantly, GTP hydrolysis is not required for membrane tethering (Ban et al., 2017), even though it is essential for membrane fusion. Finally, we show here that, in the absence of GTP, PE promotes the membrane tethering activity of Atl, the "fusion" dynamin known to rely on GTP for the membrane tethering (Kim et al., 2017; Liu et al., 2015; Pendin et al., 2011; Saini et al., 2014). It follows that the mixture of PE-enriched membranes and reduced amount of GTP in solution triggers dynamin’s functional promiscuity related to their general ability to tether membranes together and produce membrane contact sites.

Here we link the promiscuity to the ability of small Dyn2 oligomers to interact both within a single membrane and across two membranes. The former interaction fits into canonic models of Dyn2 oligomerization. It involves Dyn2 oriented in parallel and sitting in the same membrane. This lateral interaction between the Dyn2 oligomers is stimulated by GTP and results in the formation of the membrane fission machinery (Fig. 6). Conversely, membrane tethering implies dimerization between Dyn2 oligomers with anti-parallel orientation, typical for fusion dynamins (Kim et al., 2017; Pendin et al., 2011). Interestingly, some members of the dynamin superfamily, such as Atl and bacterial DLPs, are seemingly capable of both parallel and anti-parallel interactions, differentially regulated by GTP (Liu et al., 2018, 2015). Hence, weak trans-membrane tethering might be a common ancestor function of dynamins related to GTPase domains’ dimerization. Alternatively, the presence of PE in the membrane can significantly relax the orientation of Dyn2 oligomers allowing them to turn away from the membrane normal and engage laterally with the Dyn2 oligomers sitting in the opposite membrane. Indeed, the mixture of the positive membrane curvature of the small Dyn2 oligomers and the negative curvature of PE creates a morphologically labile membrane where local membrane deformations, such as membrane dimples implicated in membrane tethering and fusion (Kozlov and Chernomordik, 1998), can easily occur.

To summarize, the intrinsic curvature of small, subhelical Dyn2 oligomers provides the foundation for the diverse functionality of Dyn2. The function is adapted to lipidome and environment and is actively regulated by GTP. Under normal circumstances, the small oligomers’ curvature sensing underlies their timely accumulation in membrane fission places, such as endocytic vesicle necks. However, under specific circumstances related to the depletion of GTP and SH3 partners of Dyn2, the oligomers’ curvature activity leads to trans-membrane tethering stimulated and stabilized by conical lipid species. Through further experiments are required to evaluate this function promiscuity of Dyn2, we believe that it reflects a generic property of dynamins and might explain their deep involvement of variable membrane remodeling processes in the cell.

### Methods

#### Protein expression and purification

Dyn1, Dyn2, and Atl were expressed in Sf9 cells transiently transfected with pIEX6 constructs. Dyn1 and Dyn2 were purified by affinity chromatography using immobilized GST tagged amphiphysin-II SH3 domain as affinity ligand. The cell pellet was resuspended in buffer A (20 mM Hepes, pH 7.4, 150 mL NaCl, 1 mM DTT) and lysed by sonication. The lysate was clarified by ultracentrifugation, and after incubation, with amphiphysin-SH3 beads, the protein-bound resin was extensively washed in buffer A. Both Dyn1 and Dyn2 were eluted in buffer B (20 mM PIPES, pH 6.1-6.5, 1.2 M NaCl, 10 mM CaCl_2_, 1 mM DTT) and dialyzed overnight in buffer containing 20 mM Hepes, pH 7.2, 150mM KCl, 1 mM DTT, 1mM EDTA, 1 mM EGTA. The protein was aliquoted with 5% of glycerol, flash-frozen in liquid nitrogen, and stored at −80 °C.

Atl was purified with an N-terminal GST tag. Cell pellet was resuspended in buffer A200 (25 mM HEPES, 200 mM KCl, 10% glycerol, 2 mM DTT, 2 mM EDTA, 4% Triton- X100) and broken by sonication. The lysate was then cleared by centrifugation twice. After incubation with glutathione agarose beads the resin was first washed in A100 high detergent (25 mM HEPES, 100mM KCl, 10% glycerol, 2 mM DTT, 1 mM EDTA, 1% Triton-X100) and then in A100 low detergent (25 mM HEPES, 100 mM KCl, 10% glycerol, 2 mM DTT, 1 mM EDTA, 0.1% Triton-X100). The protein was eluted in low detergent buffer A100 by cleaving off the tag with HRV 3C protease (Pierce) overnight. The cleaved protein was aliquoted, flash-frozen in liquid nitrogen, and stored at −80 °C. Prof. Sandra Schmid generously provided aliquots containing purified Dyn2-PHDyn1. The expression and purification protocols were the same as used previously in (Liu et al., 2011). The purity of the proteins was determined by SDS-PAGE and Coomassie staining and quantified by BCA assay following the manufacturer’s instructions. Before functional experiments using purified protein, the protein was dialyzed for 2 hours in the working buffer containing 150 mM KCl, 20 mM HEPES, and 1 mM EDTA. Then, protein aggregates were removed centrifugation at 22000xg for 30 min. In the transmembrane protein Atl case, the dialysis buffer also contains 0.1% of Triton-X100 to avoid protein precipitation.

#### Preparation of LUVs

The following stocks to prepare the LUVs were used, all from Avanti Lipids: Dioleoyl-phosphatidyl-choline (DOPC), Dioleoyl-phosphatidyl- ethanolamine (DOPE), Dioleoyl-phosphatidyl-serine (DOPS), Rhodamine-DOPE (Rh-DOPE), 1,2-dipalmitoyl-sn-glycero-3-phosphoethanolamine-N-(7-nitro-2-1,3- benzoxadiazol-4-yl) (NBD-PE), cholesterol (Chol), and phosphatidylinositol 4,5- bisphosphate (PI(4,5)P_2_). Lipid mixtures containing DOPC:DOPE:DOPS:Chol:Rh-DOPE:PI(4,5)P_2_ at 38:39:10:10:1:2 or 78:0:10:10:1:2 were used, unless indicated otherwise. Where otherwise indicated, the concentration of DOPC was changed to equilibrate for changes in DOPE and/or PI(4,5)P_2_ concentrations in the lipid mixture. For the FRET-based lipid mixing experiments, we introduced the FRET pair, NBD-PE, and Rh-DOPE, at 1.5 mol% into the labeled population of LUVs, adjusting the mol% of DOPE accordingly. For all lipid mixing experiments, a population of non-fluorescence LUVs was prepared with the same lipid composition except for the fluorescence FRET pair. For Atl reconstitution, the lipid mixtures were depleted of PI(4,5)P_2_ in their composition.

All LUVs were prepared by mixing the lipid stocks in chloroform and drying first under N_2_ gas and then under vacuum for 120 min. The resulting lipid films were resuspended in the working buffer (20 mM HEPES pH 7.4, 150 mM KCl, 1 mM EDTA) to a final total lipid concentration of 1 mM. Finally, LUVs were formed by 10 freeze-thaw cycles in liquid N_2_ followed by extrusion through 100 or 400 nm pore size polycarbonate filters (Avanti Polar Lipids, USA). The desired LUVs concentrations for each experiment were achieved by dilution into the working buffer.

#### Formation of GSBs on silica and polystyrene beads from LUVs

LUVs were dialyzed in buffer containing 1 mM Hepes and 1 mM trehalose. Then 10-15 μL of the dialyzed LUVs were deposited on a Teflon film, divided into 4-5 small drops. We used 2 μL of 40 μm silica or 5 µm polystyrene beads solution (Microspheres–Nanospheres, USA) (for fluorescence-based experiments or force measurements respectively) to deposit beads over each drop and then dry them in vacuum for 20 min. For further hydration of the lipid films and formation of the GSBs, a 10 μL plastic tip was cut from the bottom to approximately 2/3 of its original size and used to take 5-6 μL of a 1M trehalose solution buffered in 1mM HEPES. The beads covered by dried lipid films were picked up and deposited into the cut tip’s trehalose solution and left to incubate for 15 min at 60°C in a humidity chamber. After incubation, the beads were transferred to the observation chamber filled with the working buffer (Velasco-Olmo et al., 2019).

Membrane tubulation experiments on GSBs were performed by adding 0.5 µM of purified proteins to the observation chamber. GSBs were incubated for 5 min at RT with the protein before imaging.

#### Binding assays

Binding efficiency between Dyn2 and LUVs was measured by pulling-down assays. LUVs prepared at 0.2 mM concentration with different DOPE percentages were incubated for 20 min with Dyn2 (0.5 μM). After incubation, the sample was centrifuged at 18000xg for 20 min to pull-down Dyn2-bound LUVs as the pellet. Dyn2 bound fraction was quantified by SDS-PAGE.*GTPase activity measurements.* The basal and assembly-stimulated GTP hydrolysis by WT and Dyn2-meGFP were measured by detecting inorganic phosphate release using a Malachite Green-based colorimetric assay (Leonard et al., 2005). Briefly, 0.5 μM Dyn2 was incubated at 37°C in the absence or presence of 400 nm LUVs (0.2 mM) in a buffer containing 20 mM HEPES (pH 7.5), 150 mM KCl, 1 mM MgCl_2_, and 1 mM GTP. Aliquots (20 μL) were drawn from the reaction mixtures after 30 min of incubation and transferred to a 96-well microplate containing 5 μL of 0.5 M EDTA to quench the hydrolysis reaction. Malachite Green stock solution (150 μL) was added to each well, and the absorbance at 650 nm was measured using a microplate reader. Free phosphate was determined from the absorbance values using a standard calibration curve.

#### NT template formation

NTs were formed in a microfluidic chamber over SU8 micropillars, as described previously (Martinez Galvez et al., 2020). Alternatively, NTs were formed by GSB rolling over a supported lipid bilayer (SLB). First, SLBs were formed by deposition of membrane lamella-covered 40 μm beads in a microfluidic channel build with PDMS over a clean (30 min sonication with EtOH) and plasma-activated (30 s of O_2_ plasma treatment) coverglass. After 10 minutes of incubation, the beads were removed from the microfluidic chamber by applying negative pressure to the channel’s outlet with a syringe pump. Then, new lamella-covered beads were added to the microfluidic device through the microfluidic channel’s inlet and rolled over the SLBs formed in the previous step. NTs were formed upon attachment to "hotspot" defects in the SLB.

#### NT radii measurement

To assess the NT radii, we first performed a calibration of the Rh-DOPE fluorescence intensity per membrane area (*ρ*0). For the calibration, we used a lipid bilayer supported on a glass coverglass prepared as previously described (Dar et al., 2015; Martinez Galvez et al., 2020). Images were acquired with Nikon Eclipse Ti-E motorized inverted microscope equipped with a CoolLed pE-4000 light source, 100x/1.49 NA oil objective, and Andor Zyla sCMOS camera. Image analysis was performed with ImageJ (Edelstein et al., 2010; Schindelin et al., 2012) and Origin software. NT radius was calculated as *rNT* = *Fl*/2*πρ0*, where *Fl* corresponds to the total Rh-DOPE fluorescence per NT length unit (Dar et al., 2015; Martinez Galvez et al., 2020).

#### Characterization of Dyn2 curvature-sensing activity

Curvature dependence of Dyn2- mEGFP adsorption (Fig. 3) was analyzed using NTs produced by pulling from GSBs as described earlier (Espadas et al., 2019). Direct comparison of Dyn2-mEGFP binding to the GSB and NT containing high amounts of PE was complicated by Dyn2-induced tubulation of GSB (see Fig. 1C). Instead, we compared the surface density of Dyn2- mEGFP fluorescence on the NTs pulled from different GSBs. The difference of the membrane tension between GSB reservoirs determined the range of the NT radii seen in Fig. 3B (Velasco-Olmo et al., 2019). The fluorescence density per a pixel of the NT length was calculated as shown in Fig. S8. The density was normalized to the maximum, curvature-independent density defined by the geometry of the Dyn2 scaffold (Fig. S2).

#### mEGFP fluorescence calibration

Borosilicate No. 1 coverslips were cleaned by sonication with pure ethanol for 30 min, and then rinsed 5 times with Milli-Q water, followed by 30-45 sec of plasma cleaning. Then, either purified mEGFP or Dyn2-mEGFP were diluted in PBS 1X to a final concentration of 1 nM and sonicated for 10 s to avoid big aggregates (Grassart et al., 2014). Images were acquired with Nikon Eclipse Ti-E motorized inverted microscope equipped with a CoolLed pE-4000 light source, 100x/1.49 NA oil objective, and Andor Zyla sCMOS camera. Low excitation power (20% of 488nm LED) was used to avoid bleaching (<10% of bleaching during 5min of continuous illumination). Images were acquired with the MicroManager software (Edelstein et al., 2010), using 2x2 binning. Icy open source image processing software (de Chaumont et al., 2021) was used to detect coverslip-bound individual mEGFP molecules and determine the position of the fluorescence peak (Fig. S11A). The background fluorescence was determined as the mean (per pixel) fluorescence measured in 4 randomly positioned ROIs containing no mEGFP molecules using imageJ (Schindelin et al., 2012). The mEGFP fluorescence was then measured as the total fluorescence integrated over 4x4 pixel ROI centered on the fluorescence peak (Fig. S11B, red square) minus the background fluorescence multiplied by 16. The resulting distribution of the mEGFP fluorescence intensity is shown in Fig. S11C. The single molecule nature of the measured fluorescence was further verified by photobleaching. Upon application of high excitation power (40% of 488 nm LED), we detected stepwise decrease of the mEFGP fluorescence (Fig. S11D, E) characteristic for single molecule photobleaching. The amplitude of the bleaching steps (46±10, SD, n=14) corresponded to the mean fluorescence intensity of mEGFP (Fig. S11C).

#### Characterization of LUV-NT tethering mediated by Dyn2

NTs doped with 0.1 mol% of 1,2-dioleoyl-sn-glycero-3-phosphoethanolamine-N-(Cyanine 5) Cy5-PE were incubated with Dyn2-mEGFP (10-20 nM) and 0.2 mM LUVs (same composition as NTs, but with 0.1 mol% Rh-DOPE as fluorophore to avoid crosstalk). Colocalized Rh-DOPE and mEGFP fluorescence spots over the NTs were selected and imaged over time with Nikon Eclipse Ti-E motorized inverted microscope equipped with a CoolLed pE-4000 light source, 100x/1.49 NA oil objective, and Andor Zyla sCMOS camera, using the MicroManager software (Edelstein et al., 2010). Further analysis of the colocalized spots was performed with ImageJ software (Schindelin et al., 2012). The frames with the highest mEGFP-fluorescence intensity in focus were background subtracted and analyzed as described above.

#### Optical tweezers force measurements of Dyn2 assembly over NTs pulled from GSBs

A counter-propagating dual-beam optical tweezers instrument equipped with light-momentum force sensors capable of direct force measurements was used in force experiments. The two lasers were brought to the same focus through opposite microscope objective lenses generating a single optical trap (Smith et al., 2003). NTs were generated in situ: pre-hydrated 5 µm polystyrene beads covered with lipid lamellas (as described earlier) were introduced into the experimental chamber containing working buffer at 22±1 °C. One 5 µm polystyrene bead was held in the optical trap and brought into contact with a 2 µm streptavidin-covered bead immobilized by suction at the micropipette tip. Then, the optical trap was moved away (and so the trapped bead) at a constant pulling speed to form a tube. Extension-shortening cycles were performed on individual tubes to check the reversibility of the force-extension curve. A constant position for the optical trap holding the lipid-covered bead was set after a suitable tether was obtained, and force was recorded in a passive mode. Dyn2 (0.5 µM) was then injected into the microfluidic chamber. Data were collected with high force (<1 pN), position (1-10 nm), and temporal (500 Hz) resolutions.

#### Sample preparation and image acquisition by CryoEM

Samples were prepared by 10-15 min at 37 °C incubation of 0.5 µM Dyn2 with 0.2 mM LUVs in the working buffer. 4 µL of the sample was placed on R2/1 Cu 300 mesh Quantifoil grids, previously hydrophilized by plasma cleaning. The sample on the grid was vitrified in liquid ethane using a Vitrobot system (FEI). Images were collected in a JEM-2200FS/CR transmission electron microscope (JEOL), operated at 200 kV and equipped with an UltraScan 4000 SP (4008x4008 px) cooled slow-scan CCD camera (GATAN). CryoEM image acquisition was performed at 400kx magnification, using a defocus range of 3-5 µm.

#### Size-quantification of LUVs-aggregates upon Dyn2 addition

0.2 mM LUVs were incubated with 0.5 µM Dyn2 for 10 min unless indicated otherwise. After incubation, the samples were transferred to a coverglass previously blocked with Bovine Serum Albumin to prevent lipid adhesion. 5 images per field of view (from 10-15 fields of view) were acquired and averaged using the Z-stack Average option in the ImageJ software. Further analysis of the area of LUVs and aggregates in pixel^2^ was performed using a constant threshold and Analyze Particles algorithm in ImageJ software (Schindelin et al., 2012).

#### Atl proteoliposome preparation

Alt was reconstituted 1:400 protein to lipid into 0.2- 0.3 mM LUVs doped with 1 mol% RhPE in the membrane, as described elsewhere (Espadas et al., 2019). The efficiency of Atl incorporation into proteoliposomes was measured by a flotation assay based on a 3–6–9–12–15–20–30–40% OptiPrep^TM^ density gradient followed by centrifugation at 45,000xg for 2 hours at 4°C. Upon centrifugation, fractions were collected from the bottom using a peristaltic pump. Rh-DOPE fluorescence of each fraction was analyzed with a 96-well plate reader, and protein content in the fractions was analyzed by SDS-PAGE (Fig. S12).

#### Lipid mixing experiments

To assay lipid mixing induced by Dyn2, a population of LUVs containing 1.5 mol% of Rh-DOPE and NBD-DOPE as the FRET pair were incubated with a population of unlabelled LUVs at 0.2 mM of each lipid mixture in the presence of 0.5 µM Dyn2 unless indicated otherwise. For Atl, 0.2 mM proteoliposomes with and without the FRET pair were mixed at a 1:1 ratio. MgCl_2_ and GTP were added as described to 5 mM and 2 mM final concentrations, respectively. The mixtures were placed in a plate reader and incubated with stirring at 37°C. NBD fluorescence (467 nm excitation/530 nm emission wavelengths) was measured over time to detect lipid mixing. Maximum NBD fluorescence was determined upon the addition of 1% TritonX-100 to the samples. This value was used to normalize the lipid mixing results, as shown in Fig. S13.

## Acknowledgments

This work was partially supported by the National Institutes of Health grant R01 GM121725 to V.A.F.; Spanish Ministry of Economy, Industry and Competitiveness grants BFU2015-70552-P to V.A.F. and A.V.S. and RYC-2014-01419 to A.V.S.; Spanish Ministry of Science, Innovation and Universities grants PGC2018-099971-B-I00 to A.V.S. and PGC2018-099341-B-I00 to B.I.; NanoMagCOST P2018 INMT-4321 and Basque Government grants IT1270-19. J.E. and A.V.O. acknowledge the FPI predoctoral fellowships from the Spanish Ministry of Economy, Industry and Competitiveness. acknowledges the predoctoral fellowships from the University of the Basque Country (UPV/EHU). IMDEA Nanociencia acknowledges support from the ’Severo Ochoa’ Program for Centers of Excellence in R&D (MINECO, Grant SEV-2016-0686).

## Supplemental Information

### Supplementary Figures

**Figure S1.**
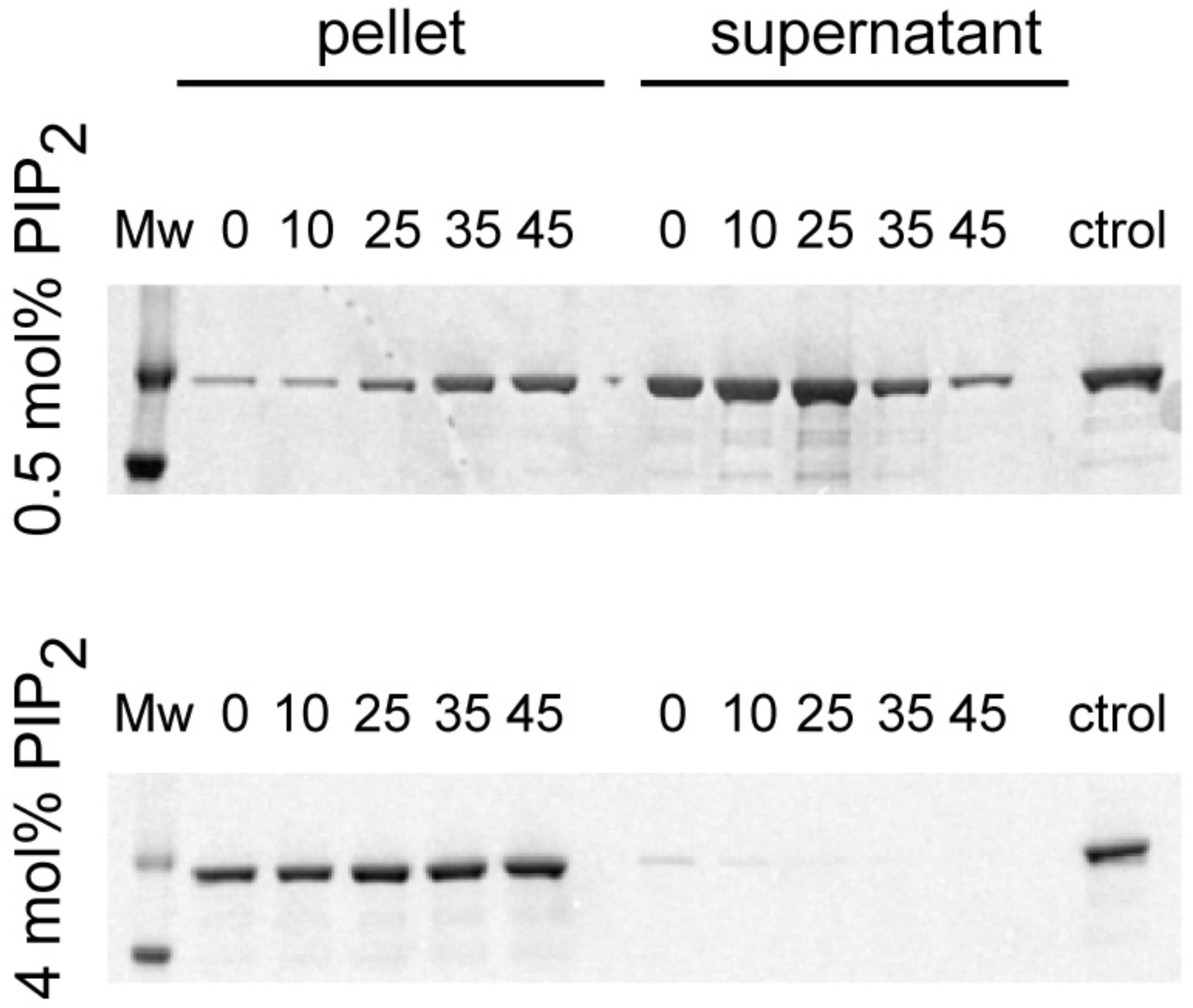
SDS-PAGE of the pellet and supernatant fractions of a pull-down assay upon incubation of Dyn2 (0.5 µM) with 400 nm LUVs (0.2 mM) with different PE (0, 10, 25, 35 and 45 mol%) and PI(4,5)P_2_ in the membrane. Upon ultracentrifugation, the protein bound to the LUVs settled in the pellet, while unbound protein was localized in the supernatant. To estimate the binding efficiency (Fig. 1A), the bands corresponding to the pellet fractions were quantified using the ImageJ algorithm and normalized against the control without LUVs.

**Figure S2.**
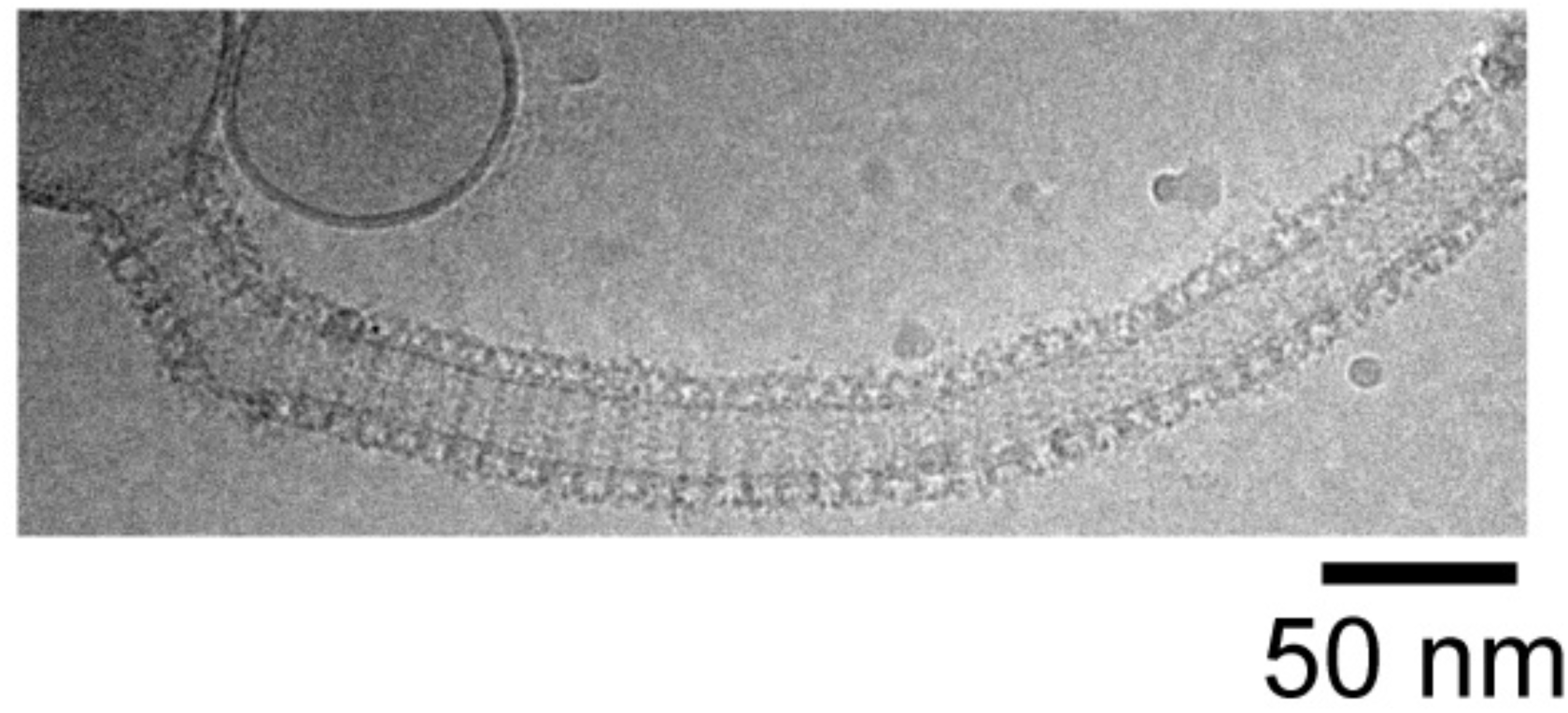
Representative cryoEM micrograph of a decorated membrane tubule formed upon incubation of 400 nm LUVs (containing 25 mol% PE and 4 mol% PI(4,5)P_2_) with 0.5 µM Dyn2 for 15 minutes.

**Figure S3.**
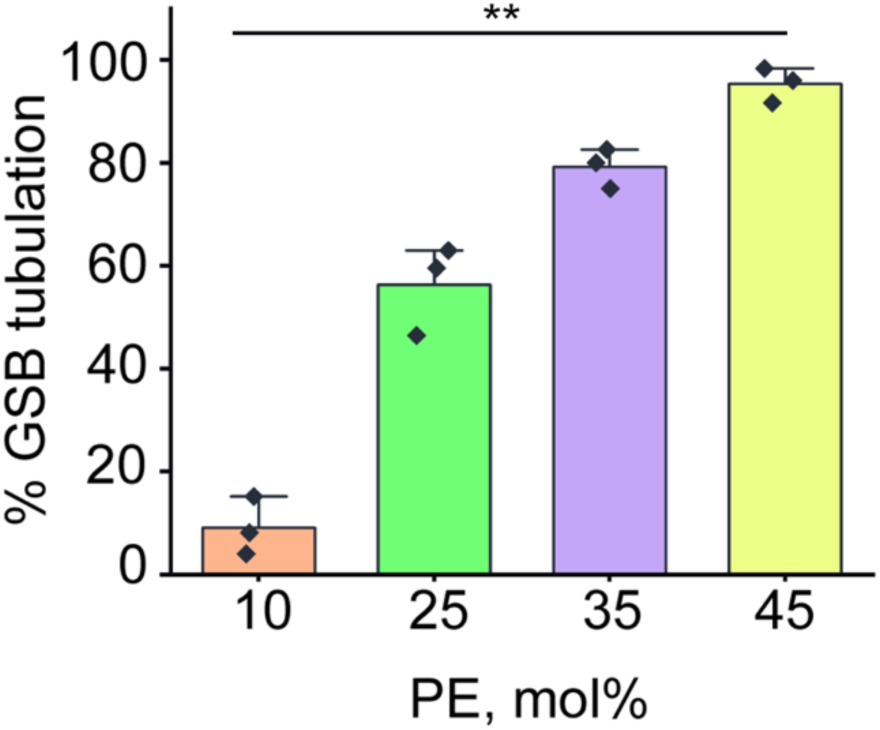
Quantification of the percentage of tubulated GSBs (see Fig. 1C) as a function of PE content in the membrane after 5 min incubation with 0.5 µM Dyn2. Error bars show SD, three independent experiments per condition. Statistical significance: one-way ANOVA with post hoc Bonferroni test, \*\**p* < 0.01.

**Figure S4.**
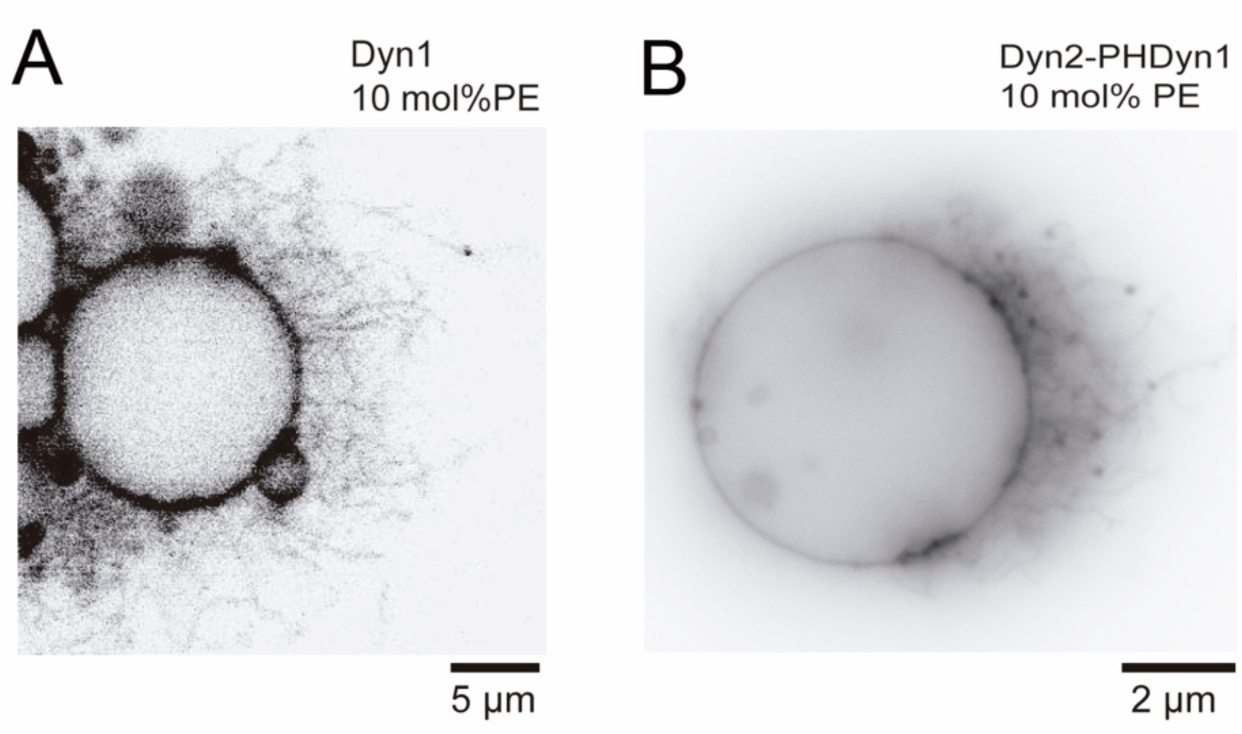
Representative images of membrane tubulation produced by Dyn1 (A) and Dyn2-PHDyn1 (B) in the GSB templates. The membranes contain 10 mol% PE and 4 mol% PI(4,5)P_2_ in the membrane. Rh-DOPE fluorescence in shown, the images were inverted for clarity.

**Figure S5.**
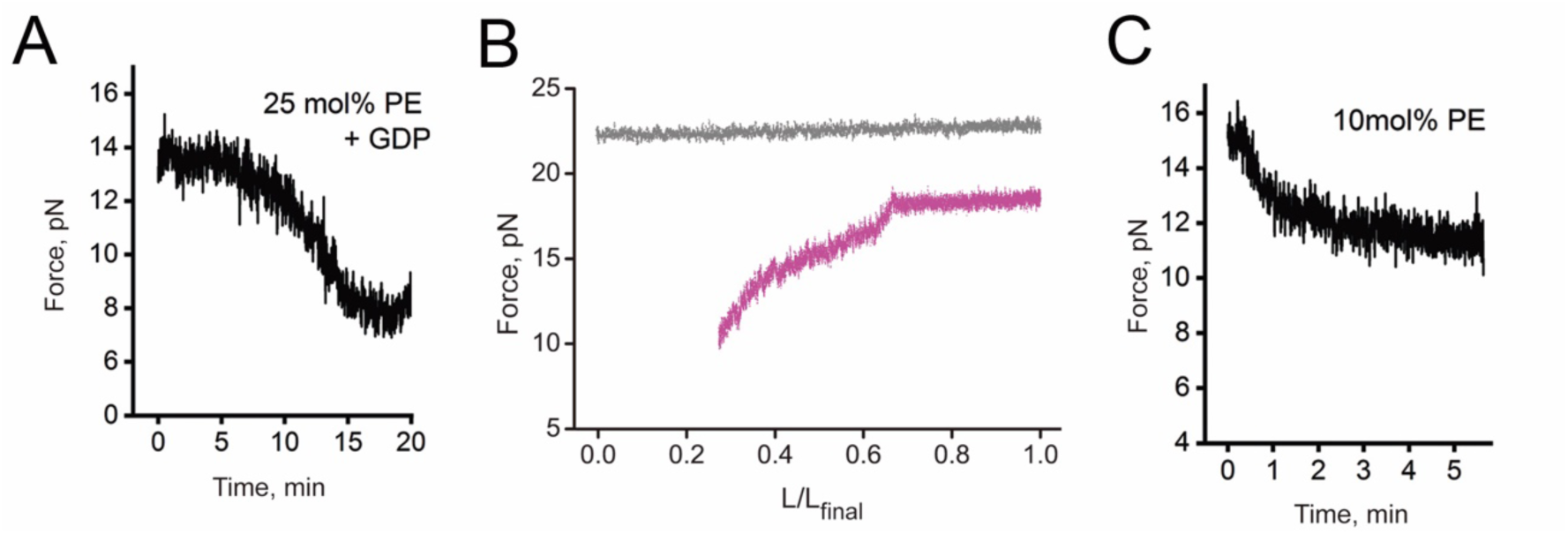
A. Force drop upon incubation of an NT (formed as in Fig. 2A from a membrane reservoir containing 25 mol% PE) with 0.5 μM Dyn2 in the presence of 1 mM GDP. A representative trace is shown. **B.** Representative traces showing the force behavior upon NT length shortening (see Fig. 2D) for a pure lipid NT (gray) and the NT in the presence of 0.5 μM Dyn2 (magenta). **C.** Force drop upon incubation of an NT containing 10 mol% PE with 0.5 μM Dyn2. A representative trace is shown.

**Figure S6.**
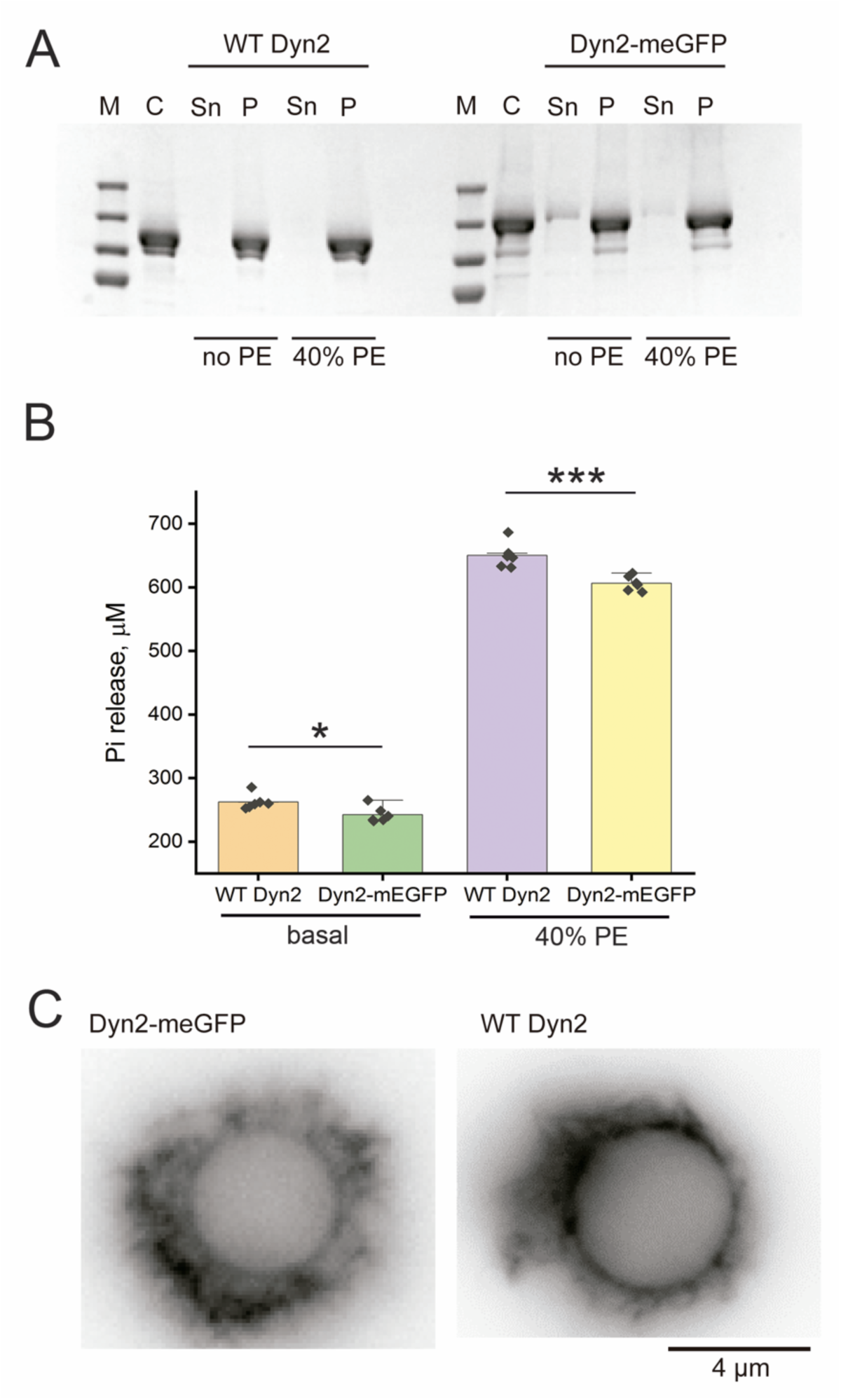
A. SDS-PAGE analysis of the pellet (P) and supernatant (Sn) fractions from pulling-down the mixtures of 0.5 μM Dyn2-WT or Dyn2-mEGFP with 0.2 mM LUVs (400 nm) after 30 min incubation. **B.** Comparison of the GTPase activity between Dyn2- WT and Dyn2-mEGFP (both at 0.5 µM) without lipid templates (basal) and in the presence of 0.2 nM 400 nm LUVs with or without 40 mol% PE (assembly-stimulated). Error bars are SD, six independent experiments. Statistical significance: two-sample unpaired *t*-test, \**p* <0.05, \*\*\**p* < 0.001. **C.** Representative images of membrane tubulation by Dyn2-mEGFP and Dyn2-WT. The images were taken 5 minutes after the protein addition, the proteinś concentration was 0.5 µM. Rh-DOPE fluorescence in shown, the images were inverted for clarity.

**Figure S7.**
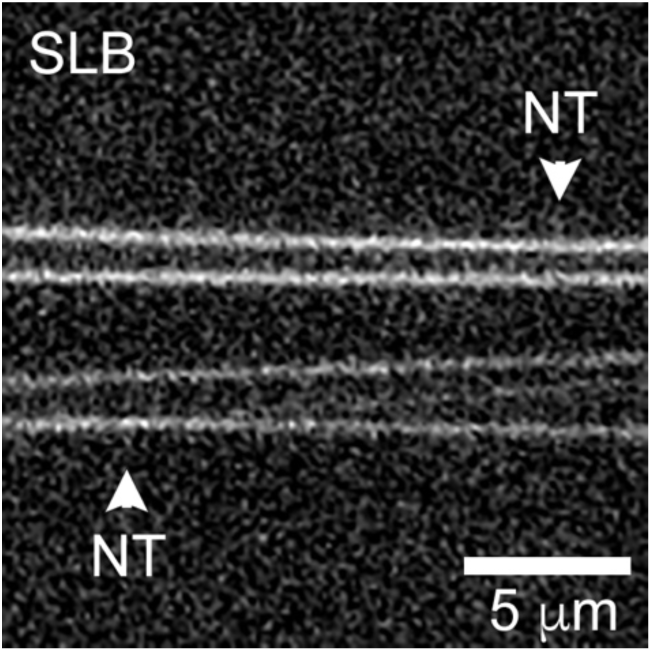
Fluorescence micrograph illustrating the formation of NTs (white arrowheads) over an SLB pre-formed on a cleansed coverglass as described in Methods.

**Figure S8.**
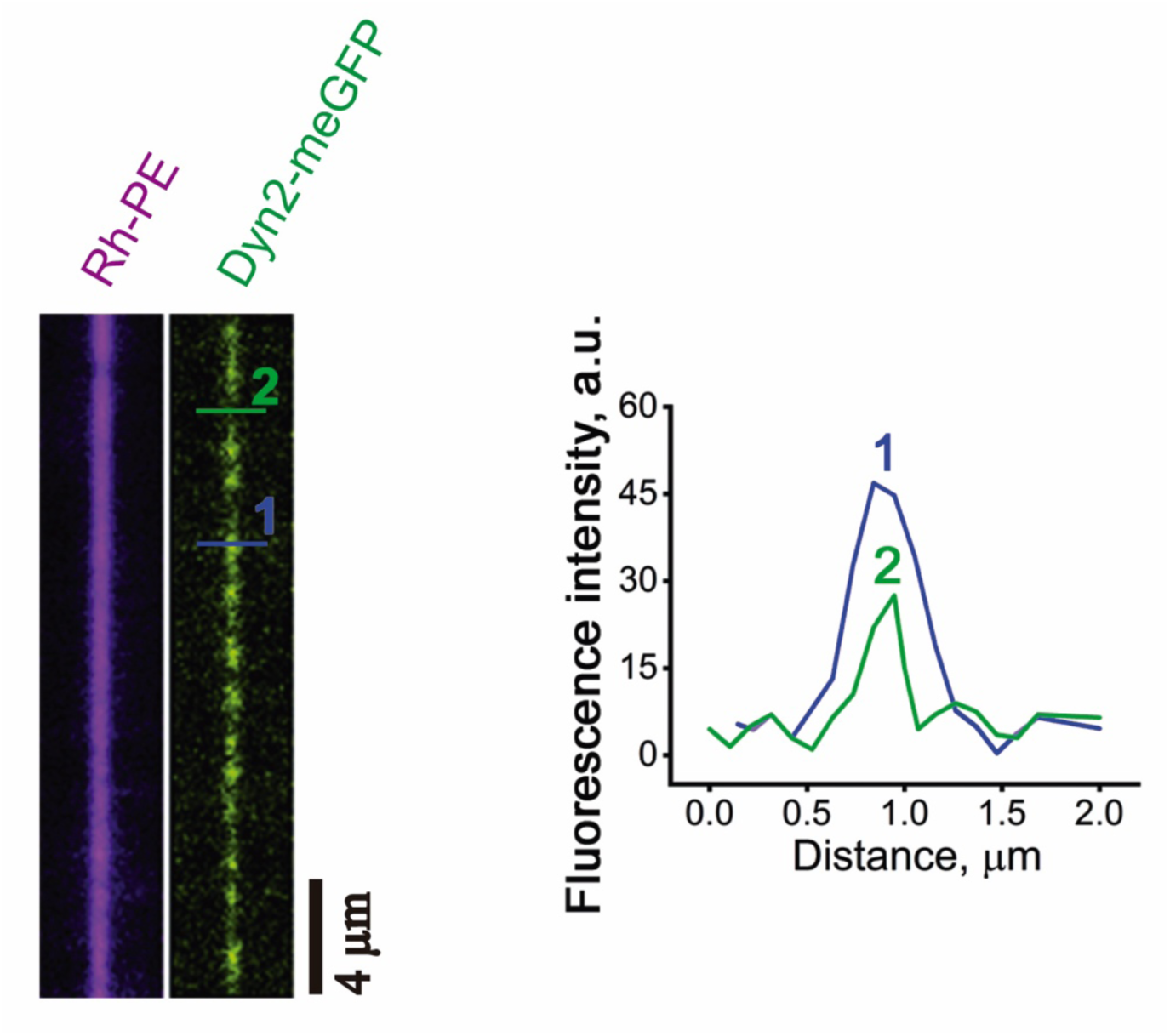
Quantification of Dyn2 helical precursors binding. The micrographs show an NT (magenta) with bound Dyn2-mEGFP (green) at different polymerization stages. The graph on the right shows the NT transversal fluorescence intensity profiles of Dyn2- mEGFP at the pre-scaffold (2, green) and complete scaffold (1, blue) stages, as shown with the corresponding lines in the micrograph on the left. The surface density of Dyn2- mEGFP over the NT at each polymerization stage was measured by normalizing the total fluorescence intensity calculated from the area under the peak in the profiles at the right against the NT lipid fluorophore integrated fluorescence intensity before Dyn2-driven membrane constriction.

**Figure S9.**
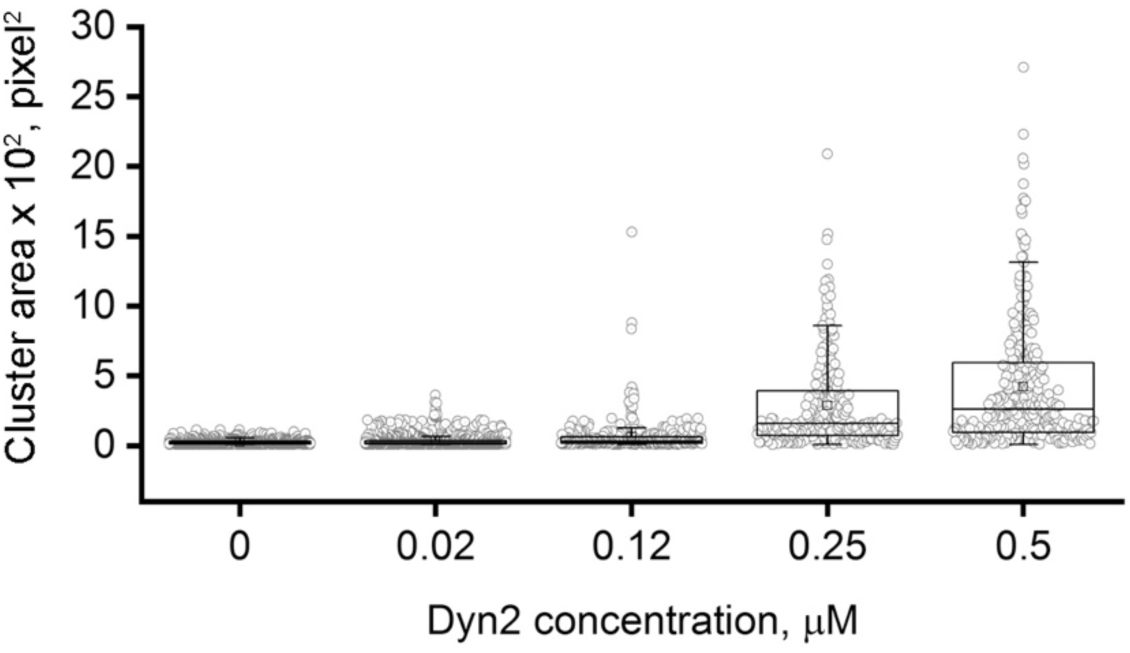
Quantification of LUV aggregation observed by fluorescence microscopy upon 30 min incubation with different Dyn2 concentrations. At least three independent experiments were done per each condition. Statistical significance: Kruskal-Wallis ANOVA test, all different at p<0.001 level.

**Figure S10.**
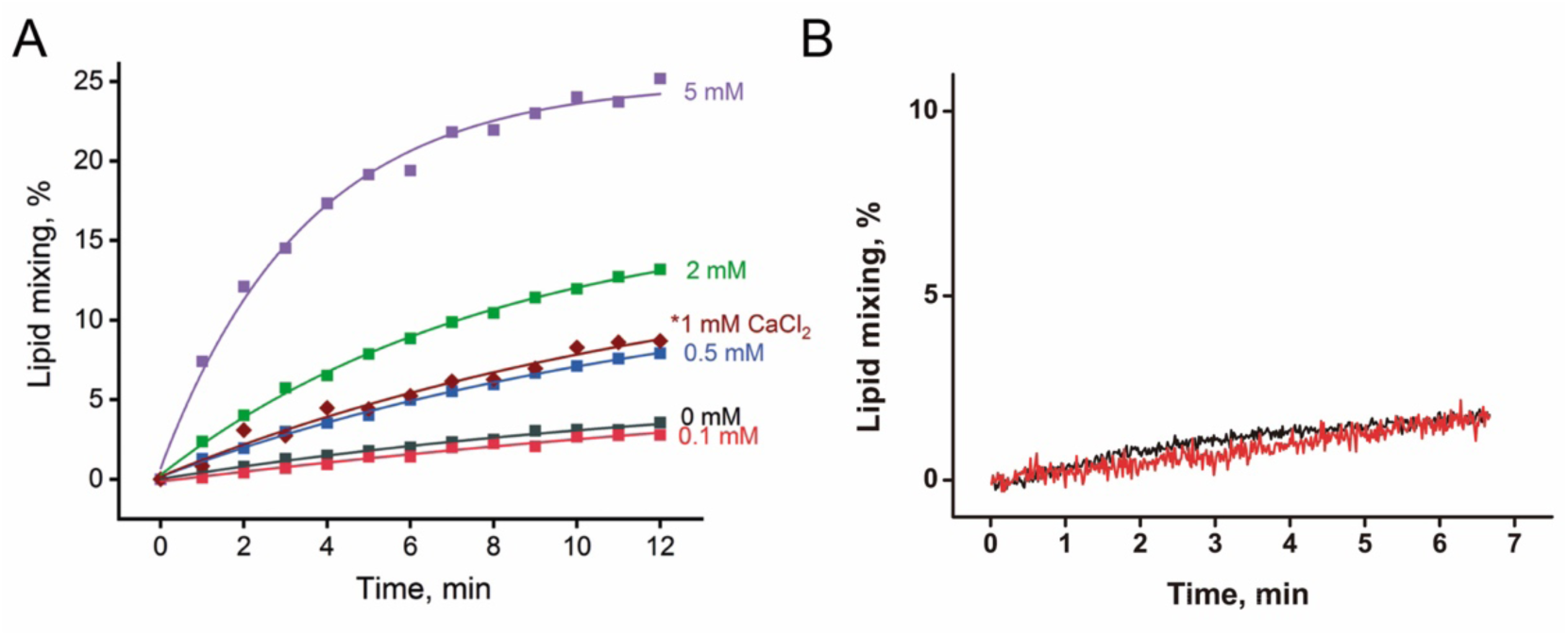
A. Effect of Mg^2+^ on lipid mixing induced by 0.5 μM Dyn2 between two populations of LUVs, as measured by FRET (see Methods). Brown curve shows the effect of the addition of Ca^2+^ instead of Mg^2+^. **B.** Effect of 5 mM MgCl_2_ (black) or CaCl_2_ (red) on lipid mixing measured as in (A), but in the absence of Dyn2.

**Figure S11.**
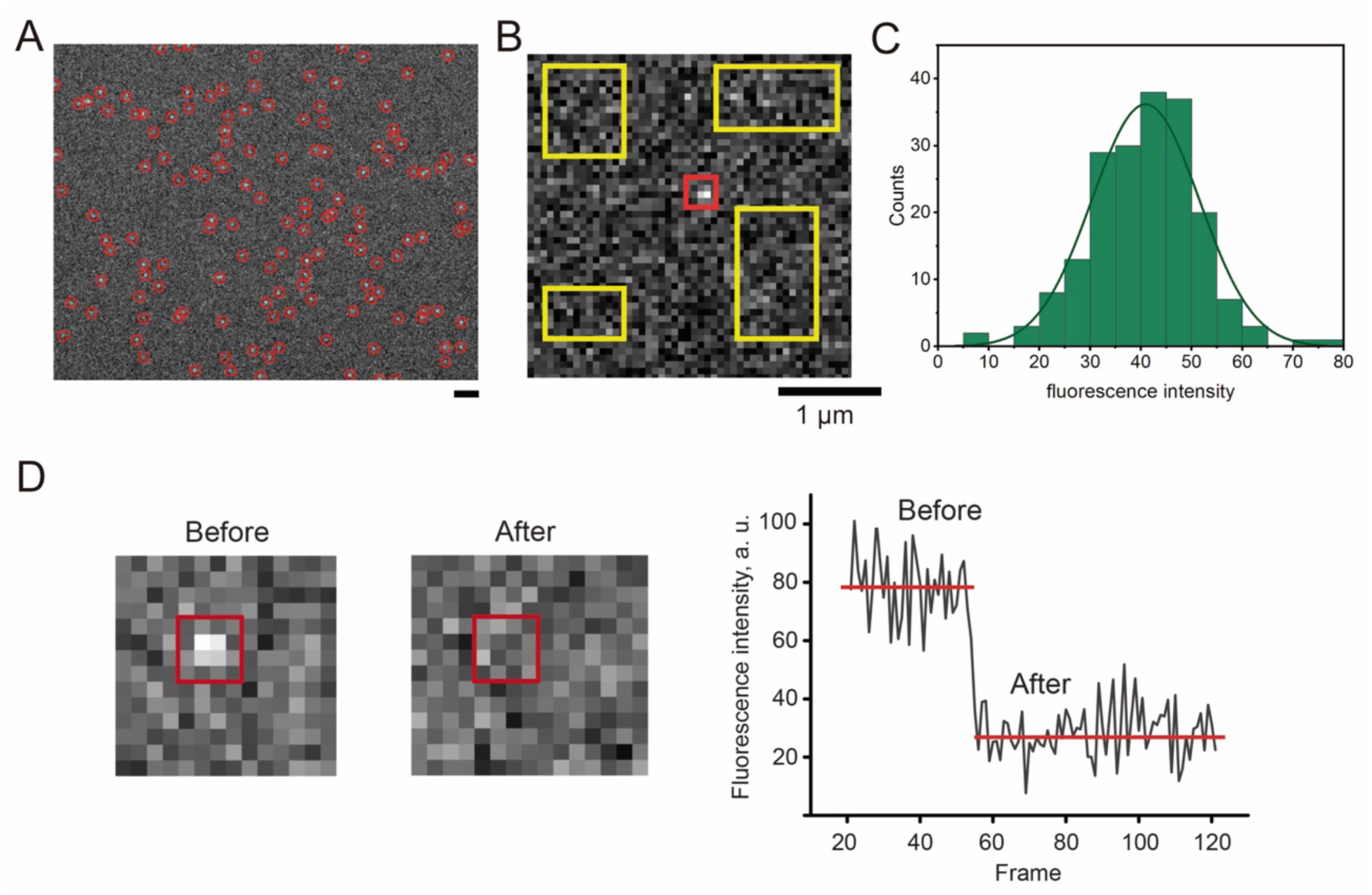
Determination of the mean fluorescence intensity of mEGFP **A.** A representative fluorescence microscopy image of the mEGFP molecules bound to the coverglass, the position of the molecules (indicated by red circles) was determined using Icy software package. **B.** Rectangular ROIs centered on the mEGFP (4x4 pixel square, red) or drawn on randomly selected areas of the coverslip free of the mEGFP (yellow) were used to calculate the mEGFP and background fluorescence. **C**. The distribution of the mEGFP fluorescence intensity. The green line shows Gaussian fit, the normality of the distribution was confirmed by Anderson-Darling test (A=0.5154). **D, E** A representative example of the stepwise photobleaching of an individual mEGFP molecule.

**Figure S12.**
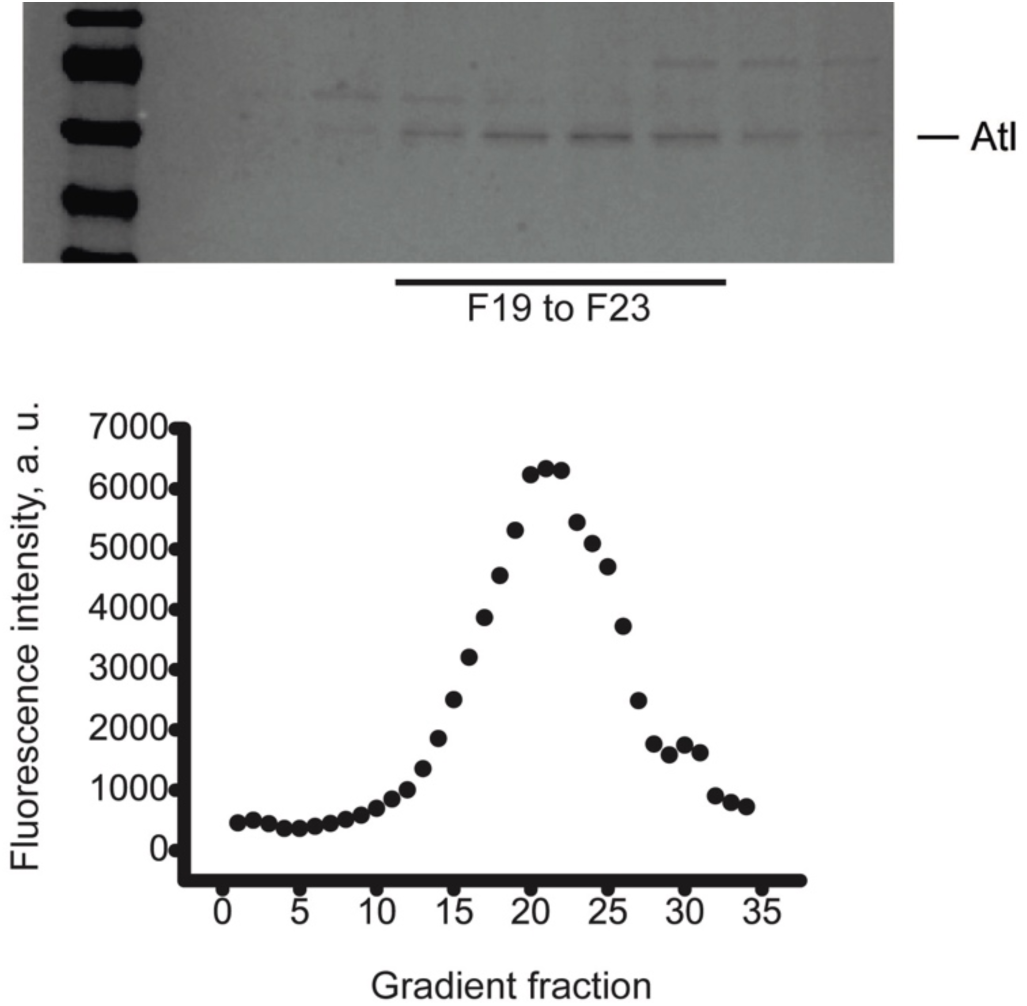
Flotation assay of reconstituted Atl proteo-LUVs in an OptiPrep density gradient. The upper SDS-PAGE shows Atl detected at each fraction of the density gradient. The fractions F19 to F23, with the maximum concentration of the protein, correspond with the highest fluorescence intensity of the lipid marker (Rh-PE, lower graph), confirming the protein’s incorporation into the proteoliposomes.

**Figure S13.**
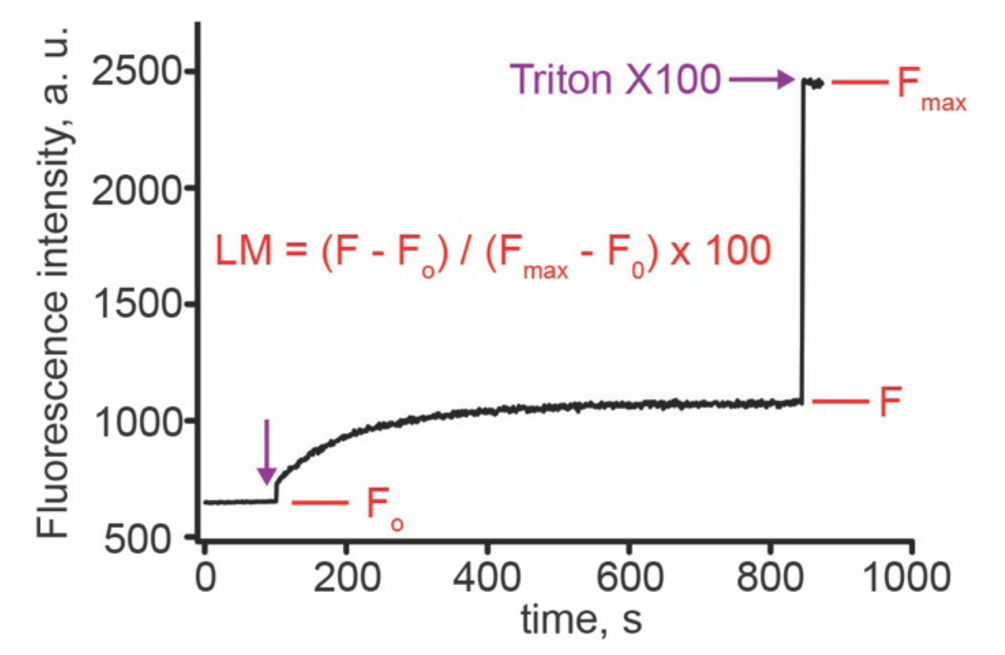
Representative curve from the FRET lipid mixing experiment. Lipid mixing endpoints were calculated by using the equation shown in red.

## Notes

### Competing Interest Statement

The authors have declared no competing interest.

